# Hetero-bivalent Nanobodies Provide Broad-spectrum Protection against SARS-CoV-2 Variants of Concern including Omicron

**DOI:** 10.1101/2022.03.08.483381

**Authors:** Huan Ma, Xinghai Zhang, Peiyi Zheng, Peter H. Dube, Weihong Zeng, Shaohong Chen, Yunru Yang, Yan Wu, Junhui Zhou, Xiaowen Hu, Yan Xiang, Huajun Zhang, Sandra Chiu, Tengchuan Jin

## Abstract

Following Delta, Omicron variant triggered a new wave of SARS-CoV-2 infection globally, adaptive evolution of the virus may not stop, the development of broad-spectrum antivirals is still urgent. We previously developed two hetero-bivalent nanobodies with potent neutralization against original WT SARS-CoV-2, termed aRBD-2-5 and aRBD-2-7, by fusing aRBD-2 with aRBD-5 or aRBD-7, respectively. Here, we resolved crystal structures of these nanobodies in complex with RBD, and found the epitope of aRBD-2 differs from that of aRBD-5, aRBD-7. aRBD-2 binds to a conserved epitope which renders its binding activity to all variants of concern (VOCs) including Omicron. Interestingly, although monovalent aRBD-5 and aRBD-7 lost binding to some variants, they effectively improved the overall affinity when transformed into the hetero-bivalent form after being fused with aRBD-2. Consistent with the high binding affinities, aRBD-2-5-Fc and aRBD-2-7-Fc exhibited ultra-potent neutralization to all five VOCs; particularly, aRBD-2-5-Fc neutralized authentic virus of Beta, Delta and Omicron with the IC_50_of 5.98∼9.65 ng/mL or 54.3∼87.6 pM. Importantly, aRBD-2-5-Fc provided *in vivo* prophylactic protection for mice against WT and mouse-adapted SARS-CoV-2, and provided full protection against Omicron in hamster model when administrated either prophylactically or therapeutically. Taken together, we found a conserved epitope on RBD, and hetero-bivalent nanobodies had increased affinity for VOCs over its monovalent form, and provided potent and broad-spectrum protection both *in vitro* and *in vivo* against all tested major variants, and potentially future emerging variants. Our strategy provides a new solution in the development of therapeutic antibodies for COVID-19 caused by newly emergent VOCs.

## Introduction

Till the beginning of 2022, the pandemic caused by SARS-CoV-2 continue to threaten global health and economic development. The widespread in global populations and adaptive evolution of SARS-CoV-2 contributed the emergence of variants of concern (VOCs), including Alpha, Beta, Gamma, Delta, and Omicron ^1-3^, which are replacing the original virus and becoming dominant strains around the world ^4, 5^. Although vaccines are considered terminator of this epidemic, the accumulation of mutations on VOCs, such as Omicron that carries 23 mutations on the S protein, has weakened the efficacy of most approved vaccines ^6, 7^. Indeed, breakthrough infections with VOCs, especially Omicron, have been reported with fully vaccinated population in many regions of the world. Coupled with the insufficient response in immunocompromised individuals to vaccines, vaccine shortages in low-income countries ^8-10^, and vaccine ineffectiveness, essential remains the development of effective prophylactic and therapeutic drugs to combat SARS-CoV-2 VOCs.

In addition to vaccines for active immunization, neutralizing antibodies for passive immunization are considered a promising alternative for the treatment of COVID-19 ^11, 12^. However, the majority of approved antibodies lose their neutralizing ability against mutant strains, particularly the Omicron variant ^13-16^. With several advantages over conventional IgG isolated from convalescent patients, variable fragments of heavy-chain-only antibodies (VHHs) derived from camelid, also called nanobodies (Nbs), are considered as an attractive alternative to traditional antibodies ^17-19^. To date, there reported a number of Nbs against SARS-CoV-2 RBD isolated from synthetic or immunized libraries ^20-41^. Nonetheless, these Nbs also face challenges with VOCs, most of them lack *in vitro* evaluation of their neutralizing activities against the currently circulating VOCs, let alone *in vivo* evaluation of protective efficacy.

Containing only one antigen binding domain, Nbs can be easily engineered into multimeric form to obtain additional binding properties ^42^. We previously developed two hetero-bivalent receptor binding domain (RBD) specific Nbs, namely aRBD-2-5 and aRBD-2-7, by tandemly fusing monovalent aRBD-2 with aRBD-5 or aRBD-7, respectively, and demonstrated potent neutralizing activities against original SARS-CoV-2 (WT) ^43^. In this study, we determined the structures of these Nbs in complex with RBD, and identified a highly conserved epitope recognized by aRBD-2. In addition, our hetero-bivalent Nbs exhibited excellent binding and neutralization to major VOCs including Alpha, Beta, Gamma, Delta, and Omicron. Especially, aRBD-2-5-Fc showed ultra-potent neutralizing activity to authentic virus of all five VOCs *in vitro*. More importantly, aRBD-2-5-Fc was able to effectively protect mice and hamsters infected with SARS-CoV-2, including the Omicron variant.

## Results

### Structural basis for potent neutralization by aRBD-2-5 and aRBD-2-7

To investigate the molecular mechanism by which aRBD-2, aRBD-5 and aRBD-7 neutralize SARS-CoV-2, a series of crystallization experiments were performed. The crystal structures of aRBD-2-7 in complex with SARS-CoV-2 RBD-tr2 (tandem repeat RBD-dimer) and aRBD-5 complexed with SARS-CoV-2 RBD were resolved at 3.2 Å and 1.8 Å, respectively. Data collection and model refinement statistics of the structures are shown in **Supplementary table 1**. Based on the structural analysis, the epitope of aRBD-2 on RBD different from that of aRBD-5 or aRBD-7, while the epitope of aRBD-5 and aRBD-7 overlaps (**Fig. 1A, B and Supplementary Fig. 1A-C**). aRBD-5 and aRBD-7 bind to the concave surface anchored by the β-hairpin of the receptor-binding motif (RBM) of RBD, while aRBD2 recognizes a different epitope closing to the lateral loop of RBM (**Fig. 1A, B, and Fig. 2A-D**).

**Fig. 1.**
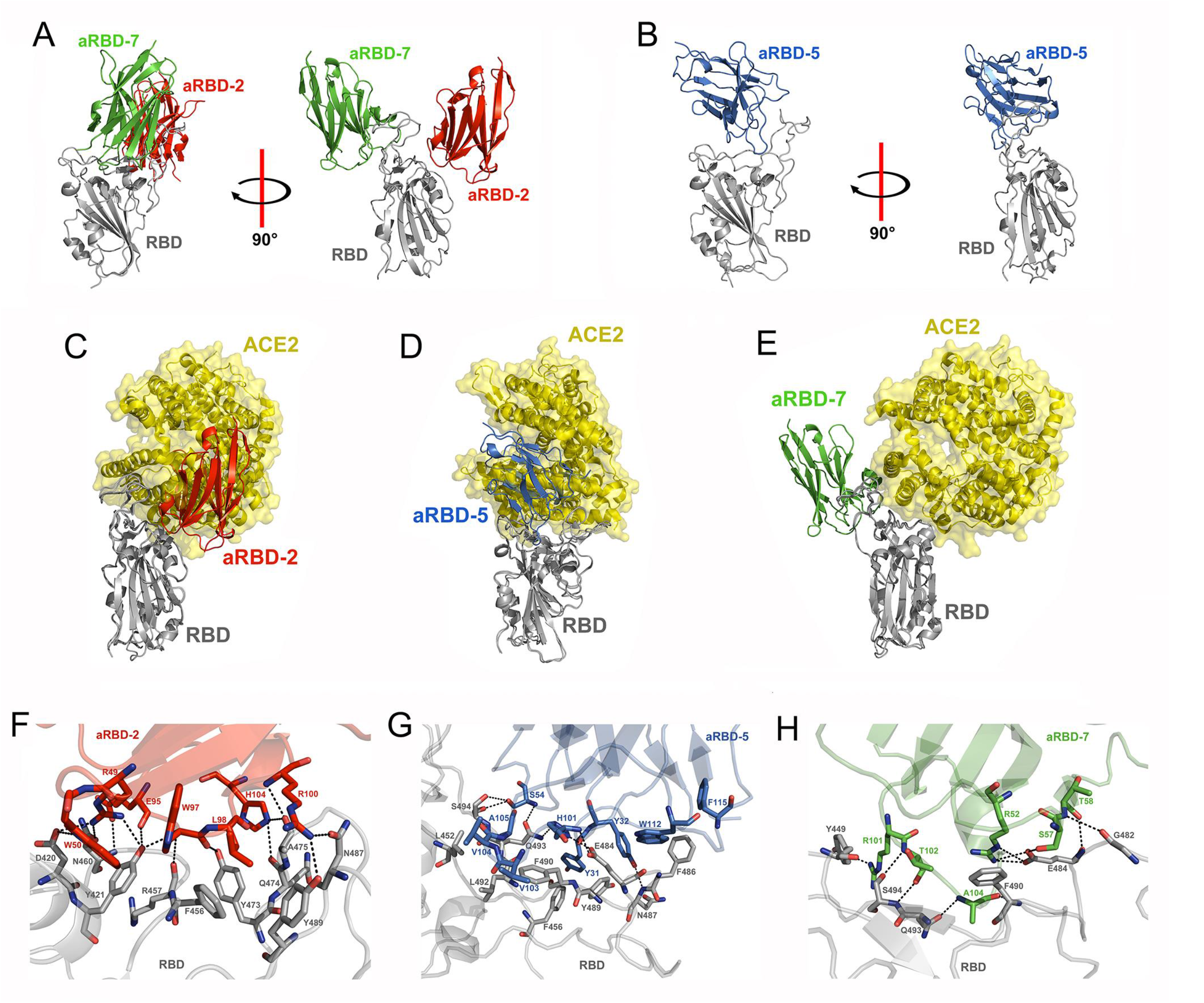
Crystal structures of aRBD-2, aRBD-5 and aRBD-7 in complex with RBD. Overall structure of aRBD-2 (red) and aRBD-7 (green) bound to RBD (grey) (A) and aRBD-5 (marine) bound to RBD (grey) (B) using cartoon presentation. The structure of aRBD-2 (C), aRBD-5 (D) and aRBD-7 (E) bound to the RBD were aligned to the crystal structure of ACE2-RBD complex (PDB: 6M0J), respectively. Zoomed-in view of the aRBD-2 (F), aRBD-5 (G) and aRBD-7 (H) with the RBD, residues that form interactions are shown as sticks, hydrogen bonds and salt bridges between Nbs and the RBD are shown as black dotted lines.

**Fig. 2.**
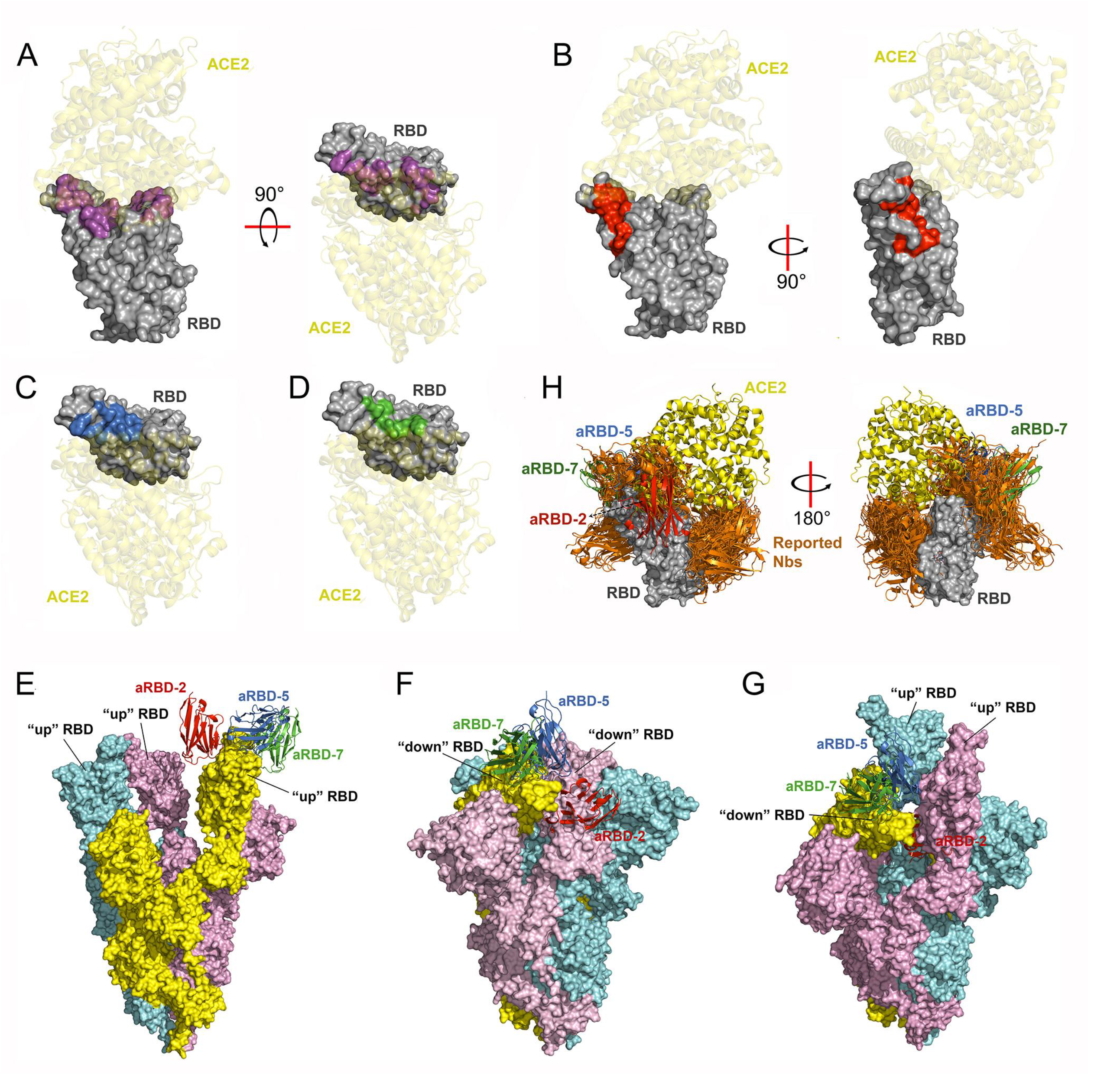
Alignment and analysis of structure of aRBD-2, aRBD-5 and aRBD-7 in complex with RBD. (A) surface display of the RBM (magenta) on RBD. (B), (C) and (D) are display of binding epitopes of aRBD-2 (red), aRBD-5 (marine), and aRBD-7 (green) on RBD, respectively. (E) Alignment of the aRBD-2, aRBD-5 and aRBD-7 in complex with the RBD to the one RBD in cryo-EM structure of the trimer spike with all “up” conformation (PDB: 7KMS). (F) Alignment of the aRBD-2, aRBD-5 and aRBD-7 in complex with the RBD to the one RBD in cryo-EM structure of the trimer spike with all “down” conformation (PDB: 7DF3). (G) Alignment of the aRBD-2, aRBD-5 and aRBD-7 in complex with the RBD to the “down” RBD in cryo-EM structure of the trimer spike with two “up” and one “down” conformation (PDB: 7KMZ). (H) Alignment the structures of aRBD-2, aRBD-5 and aRBD-7 in complex with RBD to the structures of other reported Nbs (orange) in complex with RBD deposited in PDB database, the PDB numbers of these published structures are 6ZH9, 6ZXN, 7A25, 7B27, 7C8V, 7D2Z, 7D30, 7KGK, 7KKL, 7KLW, 7KN5, 7KN6, 7KN7, 7LX5, 7MEJ, 7MFU, 7MY2, 7MY3, 7N9C, 7N9E, 7N9T, 7NKT, 7OAO, 7OAP, 7OAY, 7OLZ, 7VNB, 7KM5, 7RXD,7FG3 and 7NLL.

aRBD2 employs its CDR2 and CDR3 to bind on the RBD with an interface area of 639.1 Å2. Specifically, residues Arg49 and Trp50 of CDR2 forms three hydrogen bonds with the side chains of Asp420, Asn460 of RBD, while a salt bridge and a cation-π interaction are formed between Arg49 of CDR2 and Tyr421 of RBD, CDR3 of aRBD-2 is mainly immobilized by ten hydrogen bonds interacting with Tyr421, Agr457, Asn460, Tyr473, Gln474, Ala475, Asn487 and Tyr489 of RBD. In addition, Leu98 of CDR2 specifically interacts with Phe456 and Tyr489 of RBD through hydrophobic interactions, which further contributes to the aRBD2: RBD interaction (**Fig. 1F, Supplementary table 2**). aRBD-2 has a footprint adjacent to and obviously overlapping the ACE2 binding site, and the framework of aRBD-2 clashes with ACE2, which explains the blocking activity of aRBD-2 (**Fig. 1C** and **Fig. 2A, B**).

In contrast with aRBD-2, aRBD-7 binds to a disparate epitope on the β-hairpin of RBM (**Fig. 1A and Fig. 2B, D**). The binding of aRBD-7 to RBD is also largely contributed by CDR2 and CDR3 with a buried surface area of 669.3 Å^2^. Beside a salt-bridge (Arg52 of CDR2 to Glu484 of RBD), five hydrogen bonds were found at the interface between CDR2 of aRBD7 and RBD (**Fig. 1H, Supplemental table 2**). In addition, aRBD7 also interact with RBD through a cation-π interaction between Arg52 of CDR2 and Phe490 of the RBD. The CDR3 of aRBD-7 forms four hydrogen bonds with Phe490, Gln493 and Ser494 of RBD and a salt bridge with Tyr449 (**Fig. 1H, Supplementary table 2**). The binding epitope of aRBD-7 on RBD has a small overlap with that of ACE2, mainly formed by Tyr449 and Gln493 of RBD (**Fig. 1E, H and Fig. 2A, D**).

The binding epitope of aRBD-5 partially overlaps with that of aRBD-7. The 697.9 Å^2^ of interface area between aRBD-5 and RBD is mediated by all of its three CDRs. Except for a hydrogen bond between Tyr32 of CDR1 and Asn487 of RBD, Ser54 of CDR2 forms three hydrogen bonds with Glu493 and Ser494 of RBD, while His101 on CDR3 forms another three hydrogen bonds with Glu484 and Gln493 of RBD. Furthermore, three hydrophobic patches were formed at the interface with direct hydrophobic interactions between Tyr31 of aRBD-5 with Phe456 and Tyr489 of RBD, between Val2, Trp112 and Phe155 of aRBD-5 with Phe486 of RBD, and between Val103, Ala104 and Ala105 of aRBD-5 with Leu452, Phe490 and Leu492 of RBD, respectively. (**Fig. 1G, Supplementary table 2**). aRBD-5 stands diagonally right above the groove of the β-hairpin of RBM that interacts with ACE2, it can effectively block the interaction of ACE2 and RBD (**Fig. 1D, Fig. 2C and Supplementary Fig. 1D**).

To further analyze the neutralization mechanism of our Nbs to the intact trimeric spike protein, we superposed our crystal complex structures with spike in different conformations resolved by cryo-electron microscopy (cryo-EM). All of the three Nbs can bind to the “up” RBD in open conformation of spike protein (**Fig. 2E**), while aRBD-5 and aRBD-7 can also bind to the “down” RBD in inactive conformation of spike protein, regardless whether the adjacent RBD is “up” or “down” (**Fig. 2F, G**). Due to the steric clashes with the adjacent RBD, aRBD-2 fail to bind to the “down” RBD **(Fig. 2F, G**). Furthermore, we compared our Nb-RBD structures with all published Nb-RBD or Nb-spike structures deposited in the PDB database. Structural comparisons suggested that the epitopes of aRBD-5 and aRBD-7 overlap with some of those published Nbs, while the epitope of aRBD-2 is novel to our knowledge (**Fig. 2H, Supplementary Fig. 2**).

### Hetero-bivalent Nbs aRBD-2-5 and aRBD-2-7 bind to SARS-CoV-2 variants with high affinity

To explore the impact of the RBD mutations to our Nbs, we measured the binding affinities of aRBD-2, aRBD-5, aRBD-7 and their fusions (aRBD-2-5 and aRBD-2-7) to the RBD of WT, Alpha (B.1.1.7), Beta (B.1.351), Gamma (P.1), Delta (B.1.617.2), Delta plus (AY.1), Kappa (B.1.617.1), Lambda (C.37) and Omicron (B.1.1.529) variants, respectively. Surface Plasmon Resonance (SPR) was used with Nb-Fc fusions immobilized on the chip and RBD flowed over at different concentrations by which diminishes the avidity effects caused by the dimerization of Fc fragment. As shown in **Supplementary Fig. 3** and **Table 1**, aRBD-2 bound to Alpha, Delta, Kappa and Lambda with similar affinities as WT (1.47 nM), and to Beta, Gamma and Delta plus with slight decreased affinities, but to Omicron more than 5 fold weaker. aRBD-5 still kept the similar affinity with WT (2.3 nM) on Alpha, Delta and Delta plus RBD, but lost the binding to Beta, gamma, Kappa, Lambda and Omicron RBD. The binding of aRBD-2 and aRBD-5 to the RBD of Alpha, Delta and Delta plus are well expected, since no mutations on the epitopes. Further, the fusion protein of the two non-competitive Nbs, namely aRBD-2-5, showed a synergy effect on the affinity of binding to these RBDs, with K_D_ values of 16.8, 5.37 and <1 pM, respectively. Surprisingly, although aRBD-5 alone failed to bind to Beta, Gamma, Kappa, Lambda and Omicron RBD, the aRBD-2-5 still exhibited stronger binding affinities to these mutant RBDs than aRBD-2 alone (which is 3.28, 4.18, 1.4, 1.88 and 7.96 nM, respectively), with K_D_ values of 714, 668, 808, 53 and 171 pM, respectively. To Lambda and Omicron RBD, aRBD-2-5 has about 35-fold and 46-fold higher affinity than aRBD-2 alone.

**Table 1.**
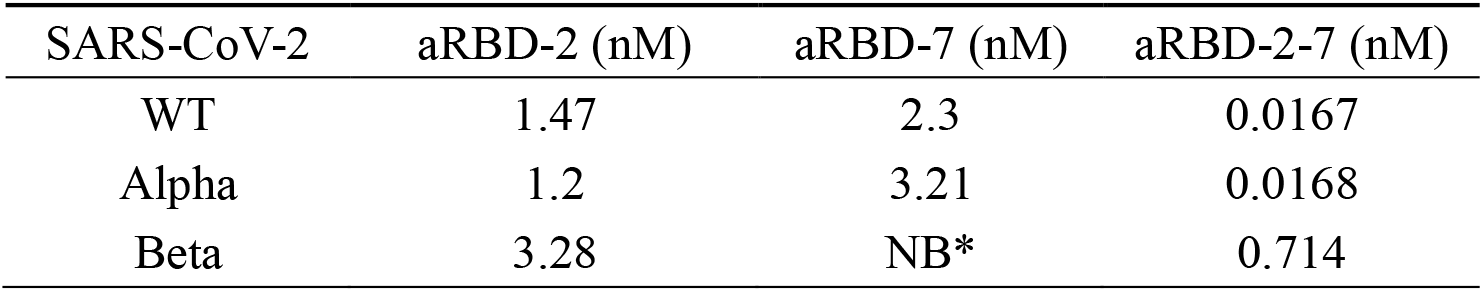
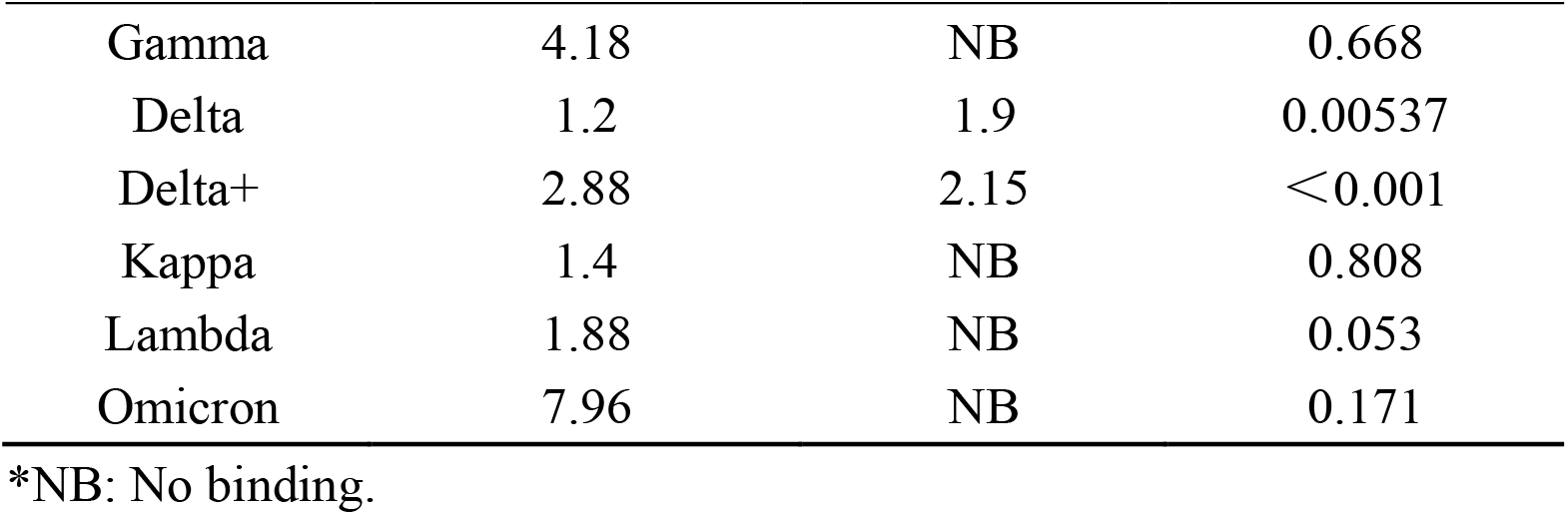
Binding affinity K_D_ of SARS-CoV-2 variant RBD to Nbs detected by SPR

Since the aRBD-7 failed to be tolerant to the acidic or alkaline solutions used for regeneration of SPR chips, ELISA was used to measure the binding of aRBD-2-7, along with aRBD-2 and aRBD-7, to the RBD of variants which were coated in the plates. Except for slightly reduced binding to the RBD of Beta (EC_50_ of 4.49 nM) and Omicron (EC_50_ of 3.24 nM) compared to WT (EC_50_ of 1.15 nM), aRBD-2-Fc showed similar binding activity to the other six variants (**supplementary Fig. 4** and **Table 2**), these results are roughly consistent with that of SPR. aRBD-7-Fc bound to the WT and Alpha RBD with similar EC_50_ of 0.117 nM and 0.141 nM, respectively, but completely or partially lost the binding to the other seven variants. As in the case of aRBD-2-5-Fc, aRBD-2-7-Fc displayed much higher binding to all mutant RBDs over aRBD-2-Fc alone (**supplementary Fig. 4** and **Table 2**). Taken together, aRBD-2 alone maintains the binding activity to the RBD of all tested variants, and fusing nanobodies with non-overlapping epitopes, such as aRBD-5 or aRBD-7, to aRBD-2 can effectively improve the overall affinity to these variants.

**Table 2.** Binding EC_50_ of Nb-Fc fusions to SARS-CoV-2 variant RBD detected by ELISA

| SARS-CoV-2 | aRBD-2 (nM) | aRBD-7 (nM) | aRBD-2-7 (nM) |
| --- | --- | --- | --- |
| WT | 1.146 | 0.117 | 0.156 |
| Alpha | 1.204 | 0.141 | 0.191 |
| Beta | 4.490 | NB* | 0.517 |
| Gamma | 1.834 | NB | 0.135 |
| Delta | 1.537 | ND<sup>#</sup> | 0.149 |
| Delta+ | 1.330 | ND | 0.116 |
| Kappa | 1.000 | NB | 0.093 |
| Lambda | 1.953 | NB | 0.165 |
| Omicron | 3.238 | NB | 0.179 |

\*NB: No binding. <sup>#</sup>ND: Not defined.

### Hetero-bivalent Nbs exhibit potent neutralizing ability against SARS-CoV-2 variants *in vitro*

Using Fc fusion to form homodimer can increase the avidity and half-life of Nbs, so Nb-Fc fusions were further tested for neutralization. We measured the neutralizing activity of the aRBD-2-5-Fc and aRBD-2-7-Fc on cellular entry of the SARS-CoV-2 variants including Alpha, Gamma, and Kappa using a SARS-CoV-2 micro-neutralization assay (**Supplementary Fig. 5**). As a control, Nb21-Fc, a previously described nanobody with strong neutralizing activity against the WT virus ^37^, lost the neutralizing activity to Gamma and Kappa variant, but retains the activity to Alpha with IC_50_ of 7.58 ng/mL. To the contrast, aRBD-2-5-Fc can block all of the three tested variants, including Alpha, Gamma, and Kappa variants, with IC_50_ of 5.63, 11.97 and 8.47 ng/mL, respectively. The IC_50_ of aRBD-2-5-Fc for Alpha was slightly lower than or close to that of Nb21-Fc. Comparing to aRBD-2-5-Fc, aRBD-2-7-Fc neutralizes Alpha with similar activity, but 24 times stronger for Gamma while 2.5 times weaker for Kappa (**supplementary Fig. 5A-D**).

We also tested the neutralizing activity of the Nb-Fc fusions to WT, Beta, Delta and Omicron variants using plaque reduction neutralization assay (PRNT) (**Fig. 3**). In addition to the Nb21-Fc, another FDA approved conventional IgG type antibody, Sotrovimab (S309 ^44^), was also used as a control. In this assay, Nb21-Fc lost the neutralizing activity to Beta and Omicron variants, but retains the activity to WT and Delta with IC_50_ of 8.95 and 4.90 ng/mL, respectively (**Fig. 3**). Both aRBD-2-5-Fc and aRBD-2-7-Fc potently neutralized WT, Beta, Delta and Omicron variant with IC50 from 5.98∼28.95 ng/mL. The neutralizing activities of aRBD-2-5-Fc, aRBD-2-7-Fc and Nb21-Fc against WT and Delta were roughly similar (within a 2.5-fold difference), while neutralization of aRBD-2-5-Fc and aRBD-2-7-Fc were much stronger than that of Sotrovimab, especially for Omicron, aRBD-2-5-Fc and aRBD-2-7-Fc exhibited ∼123 and 27-folds higher potency than Sotrovimab, respectively (**Fig. 3**). Taken together, these results showed that our hetero-bivalent Nbs can neutralize the cell invasion of SARS-CoV-2 VOCs with ultra-high potency.

**Fig. 3.**
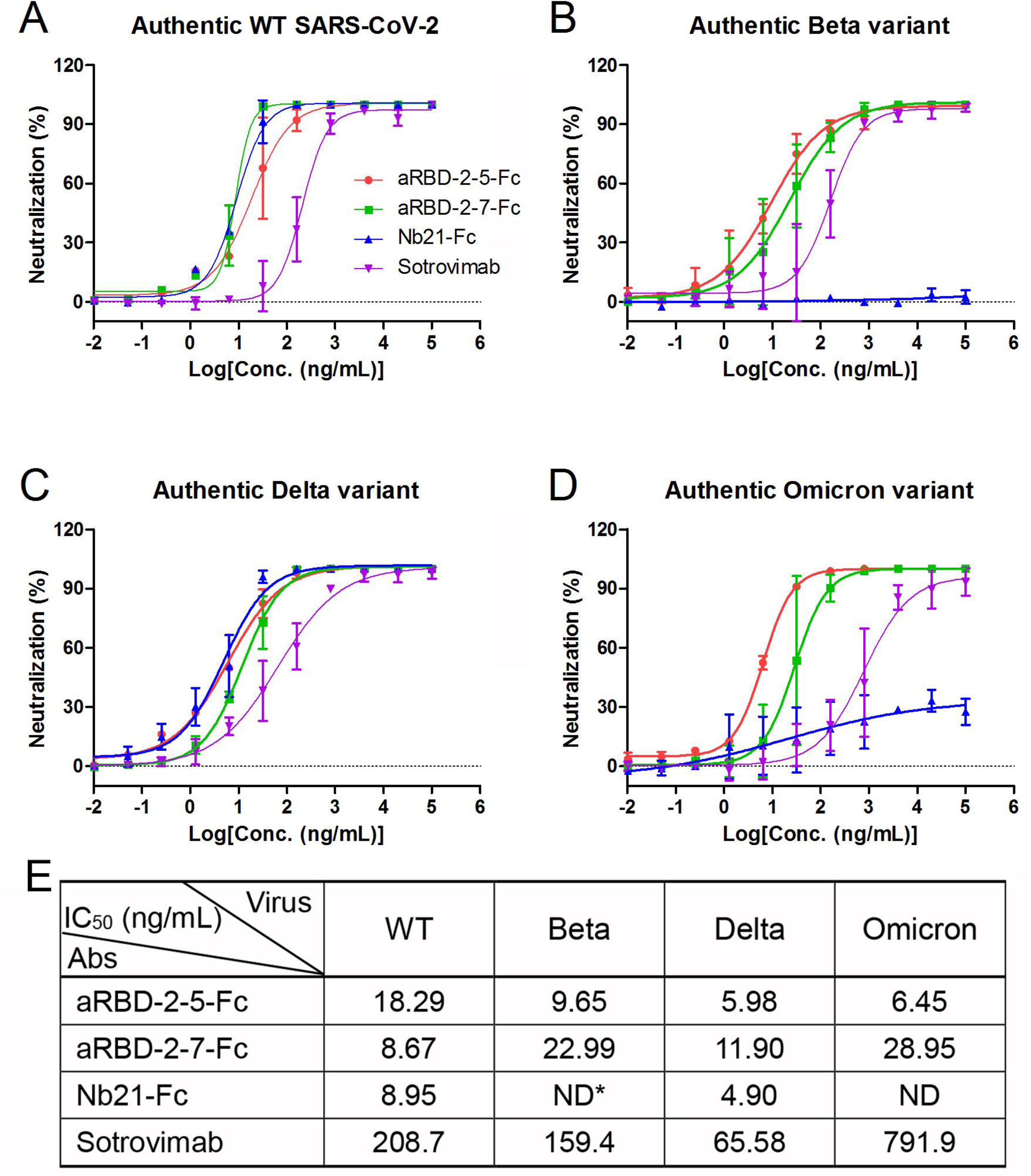
*In vitro* neutralization to SARS-CoV-2 variants by aRBD-2-5-Fc and aRBD-2-7-Fc using PRNT assay. The serially diluted antibodies were incubated with ∼75 PFU of authentic WT(A), Beta(B), Delta (C) and Omicron (D) variants, respectively. The mixture was then added to Vero E6 cells in 24-well plates. After 3 days of infection, the plates were fixed with 8% paraformaldehyde and stained with 0.5% crystal violet, and plaques were revealed by washing with distilled water and subsequently enumerated. The data was calculated by fitting the inhibition from serially diluted antibody to a sigmoidal dose-response curve. Error bars indicate the mean ± SD from two replicates. (E) is the summary of neutralizing IC_50_. *ND means neutralizing activity not detected.

### aRBD-2-5-Fc protects mice from infection of WT and mouse-adapted SARS-CoV-2

aRBD-2-5-Fc showed ultra-potent neutralizing activity at cellular level, we sought to investigate its efficacy against SARS-CoV-2 infection in animal models. aRBD-2-5-Fc intraperitoneally (i.p.) administered at 10 mg/kg dose effectively protected weight loss and death of K-18 hACE2 transgenic mice infected intranasally (i.n.) with 2 ×10^4^ PFU of WT SARS-CoV-2. (**Fig. 4B, C**). Furthermore, aRBD-2-5-Fc can provide protection from weight loss and death to A/J mice intratracheally infected with 1 ×10^5^ PFU of mouse-adapted SARS-CoV-2 virus MA10 ^45^ at the dose as low as 1 mg/kg (**Fig. 4D, E**). The mouse-adapted MA10 virus bears the RBD containing Q493K, Q498Y and P499T mutations, among which Q493 is one of the binding sites of aRBD-5. This result again demonstrates the resistance of aRBD-2-5-Fc to mutation escape.

**Fig. 4.**
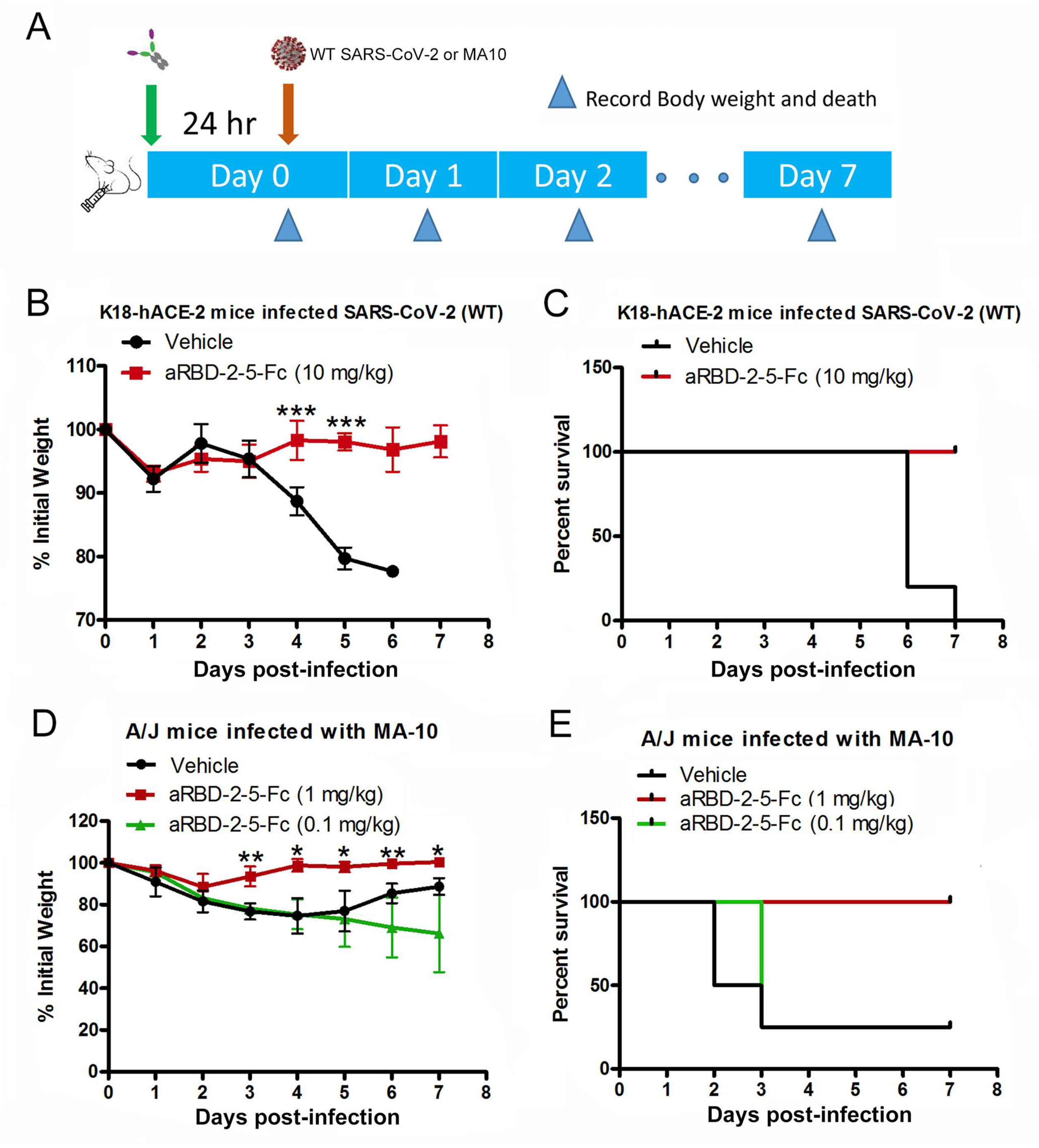
Protective efficacy of aRBD-2-5-Fc in mice models infected with SARS-CoV-2. (A) Animal experiment scheme. (B) and (C) is the body weight and death of Vehicle/aRBD-2-5-Fc protected K18-hACE2 transgene mice infected (i.n.) with 2 ×10^4^ PFU of original SARS-CoV-2, respectively. Five mice per group. (C) and (D) is the body weight and death of Vehicle/aRBD-2-5-Fc protected A/J mice infected (intratracheally) with 1 ×10^5^ PFU of mouse-adapted SARS-CoV-2 MA10, respectively. Four mice per group. Error bars indicate mean ±SD.

### aRBD-2-5-Fc provides hamsters prophylactic and therapeutic protection against Omicron variant

Omicron containing multiple number of mutations is spreading globally, so we further evaluated the prophylactic and therapeutic efficacy of aRBD-2-5-Fc against Omicron variant in hamster model. Prophylactic group of hamsters were administered with one dose of aRBD-2-5-Fc (10 mg/kg, i.p.) 24 hours (hr) before challenge (1 ×10^4^ PFU, i.n), and therapeutic group of hamsters were administered with one dose of aRBD-2-5-Fc (10 mg/kg, i.p.) 3 hr after challenge (1 ×10^4^ PFU, i.n). Four days post infection (d.p.i.), the animals were sacrificed, and trachea and lung tissues were collected to determine the viral RNA load and infectious virus particles **(Fig. 5A)**. Overall, mean levels of viral RNA in the trachea and lungs were reduced in both prophylactic and therapeutic hamsters compared to controls (**Fig. 5B**). Although viral RNA was still detected in the trachea and lungs of the hamsters in the prophylactic and therapeutic group, viral titers in these tissues were under the limit of detection (**Fig. 5C**). In contrast, infectious virus was detected in these tissues in control group (**Fig. 5C**). We also monitored the body weight of the hamsters, but since Omicron only causes attenuated disease in hamsters^46^, no significant differences were observed between the three groups (**supplementary Fig. 6**).

**Fig. 5.**
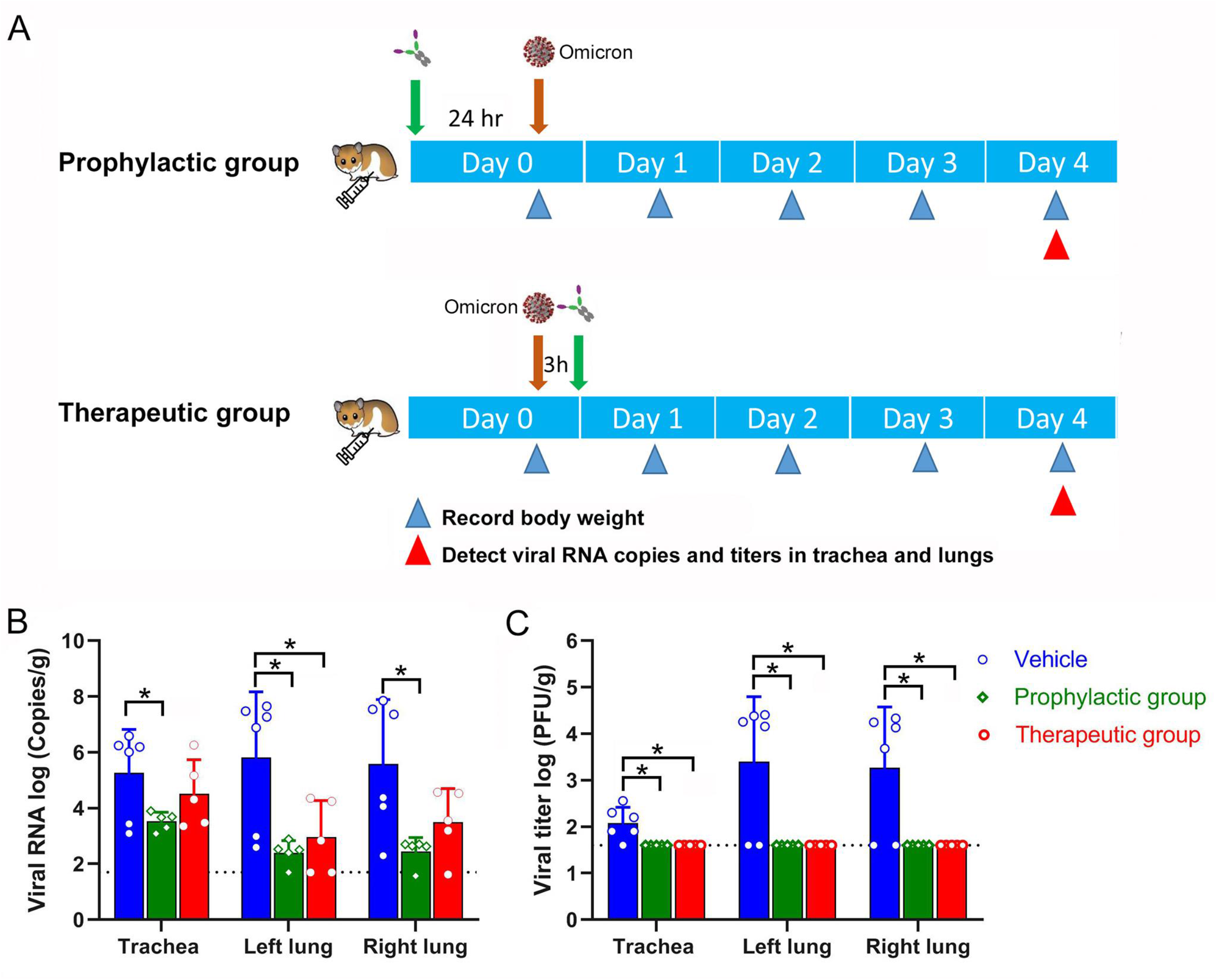
Prophylactic and therapeutic efficacy of aRBD-2-5-Fc in hamsters infected with Omicron variant. (A) Aminal experiment scheme. (B) Viral RNA (log_10_(RNA copies per g)) detected in trachea, left lung and right lung of hamsters challenged with Omicron virus at 4 d.p.i.. (B) Titers of virus (log_10_(RNA copies per g)) in trachea, left lung and right lung of hamsters challenged with Omicron virus at 4 d.p.i. measured by plaque assay. Error bars indicate mean +SD.

## Discussion

Escape mutations on the RBD of emerging SARS-CoV-2 variants have posed a tremendous challenge for therapeutic antibody and vaccine development, therefore, we further characterize the efficacy of our previously developed hetero-bivalent Nb tandems (aRBD-2-5 and aRBD-2-7) to SARS-CoV-2 variants. We first determined the crystal structures of aRBD-2, aRBD-5, and aRBD-7 that make up the hetero-bivalent tandems in complex with RBD. aRBD-2 interacts with D420, Y421, F456, R457, N460, Y473, Q474, A475, N487 and Y489 on RBD, which are highly conserved, without any mutation in current VOCs (Alpha, Beta, Gamma, Delta, and Omicron) and three additional variants of interest (VOIs) (Delta plus, Kappa, and Lambda). Except for K417 and S477 located at the edge of the aRBD-2 interface, the other mutation sites are far away from the interface (**Supplementary Fig. 7A**). This is consistent with the results of od binding assays, that is, except for Beta, Gamma, Delta Plus RBD that contain K417 mutation and Omicron RBD contains K417 and S477 mutations show a slightly weakened affinity to aRBD-2, the affinity for other mutant RBDs were not affected (**Table 1**). D420 and Y421 are highly conserved in multiple isolated sarbecovirus ^47-53^, while F456, A475, N487 and Y489 are ACE2 contacting residues on RBD ^54^, these 6 sites are not likely to be mutated. More interestingly, further structural alignment revealed that the epitope of aRBD-2 is different from that of the other reported Nbs in PDB database, and aRBD-2 does not clash with most of these Nbs upon binding to RBD. Therefore, we conclude that aRBD-2 is a mutation-tolerant nanobody targeting a novel epitope on RBD.

Both aRBD-5 and aRBD-7 interact with the E484 and Q493 of RBD, which explains why both of them lost the binding to the E484 mutated RBDs contained in the Beta, Gamma, Kappa and Omicron variants. The E484 mutation is notorious, it causes not only steric clashes but also a charge reversion at antibody binding interface, which decreases neutralization of polyclonal convalescent plasma by as much as >100-fold in some individuals ^55^. Many potent neutralizing monoclonal antibodies, such as 2-15, LY-CoV555, C121 and REGN10933, completely abolished or dramatically reduced neutralization activity by the E484 mutation^56^. The activity of a high-affinity (K_D_ less than 1 pM) nanobody Nb21 used as a control in this study was also abolished by the E484 mutation. Other Nbs, such as H11-H4^22^, TY1^23^, VHH E^26^, C5, H3^31^, Re5D05^32^ and so on, that directly bind to the receptor binding motifs may also be greatly weakened by the E484 mutation. The abolishment of binding between aRBD-5 and Lambda may be due to the loss of hydrophobic interaction caused by L452Q and F490S (**Supplementary Fig. 7B**). L452R may also slightly affect the binding of aRBD-5 to Delta and Delta Plus. F490 of RBD is recognized by aRBD-7, it should be the reason why aRBD-7 failed to bind to Lambda RBD (**Supplementary Fig. 7C**). The longer side chain formed by the mutation of L452R located at the edge of the aRBD-7 interface may cause aRBD-7 to be pushed away, which may be the reason why aRBD-7 failed to bind Delta and Delta plus (**Supplementary Fig. 7C**).

SPR is the gold-standard for measurement of antibody and antigen interactions. Using this technique, we found that although aRBD-5 itself does not bind Beta, Gamma, Kappa, Lambda and Omicron RBD, it still contributes to the increased binding of aRBD-2-5 to these mutant RBDs. Compared with the affinity of aRBD-2, the affinity of aRBD-2-5 to Beta, Gamma, Kappa, Lambda and Omicron RBD is 4.6, 6.3, 1.7, 35.5 and 46.5 times higher, respectively (**Table 1**). Similar results were also observed in the binding assay of aRBD-2-7 by ELISA (**Table 2**). A possible explanation is that fusion with aRBD-5 or aRBD-7 facilitates aRBD-2 close to the RBD which enables it to bind to the non-mutated sites. In brief summary, our results highlight the importance of preparing hetero-bivalent tandem Nbs to overcome the escape of mutant strains.

A flexible linker composed of three repeats of GGGGS was employed for fusing aRBD-2 with aRBD-5 or aRBD-7 in a “tail to head” form to construct aRBD-2-5 or aRBD-2-7 hetero-bivalent nanobody, the length of the 15-amino acid linker is about 54 Å when in an extended form. We aligned our Nbs to Cry-EM structure of trimeric spike based on our complex structures, and further measured the distance between the C-terminus of aRBD-2 and the N-terminus of aRBD-5 or aRBD-7. The observed distance between aRBD-2 and aRBD-5 on the same “up” RBD (26.2 Å, **Supplementary Fig. 8A**) or two different “up” RBD (42.4 Å, **Supplementary Fig. 8B**) is less than the length of the 3(GGGGS) linker, thus aRBD-2 and aRBD-5 in aRBD-2-5 tandem can simultaneously bind to the same RBD or two different “up” RBDs on spike. While aRBD-2 and aRBD-7 in aRBD-2-7 was neither allowed to simultaneously bind the same RBD (the distance is 64.3 Å, **Supplementary Fig. 8C**), nor respectively two “up” RBD (the distance is 80.8 Å, **Supplementary Fig. 8D**), the only feasible binding mode to warrant the aRBD-2 and aRBD-7 in aRBD-2-7 tandem simultaneously bind to the spike is that aRBD-2 binds to an “up” RBD, and aRBD-7 binds to the adjacent “down” RBD (the distance is 47.4 Å, **Supplementary Fig. 8E**), this binding mode of aRBD-2-7 would lock the “down” RBD in the closed state.

Encouraged by the high affinity of aRBD-2-5 and aRBD-2-7 for binding to RBD, we further examined their neutralizing activity against authentic VOCs *in vitro*. The results showed that both aRBD-2-5-Fc and aRBD-2-7-Fc have potent neutralizing activities against all five VOCs tested in two different neutralization assays, their IC_50_ were single-digit ng/mL to the ten-digit ng/mL level, and their neutralization activities toward the Beta, Delta and Omicron strains are much stronger than the approved antibody Sotrovimab (**Fig. 3**), a traditional IgG antibody targeting conserved epitopes on sarbecovirus. Based on such excellent neutralizing activity, evaluation on prophylactic and treatment protection in mice or Syrian golden hamster animal models were carried out. At a dose of 10 mg/kg, aRBD-2-5-Fc successfully prevented weight loss and death in K18-hACE2 transgenic mice infected with the WT SARS2-CoV-2 (**Fig. 4B, C**). Even at a low dose of 1 mg/mL, aRBD-2-5-Fc still exhibited effective protection on weight loss and death of A/J mice infected with mouse-adapted SARS-CoV-2 that contains Q493K, Q498Y and P499T mutations on RBD (**Fig. 4D, E**). Finally, we further tested the prophylactic and therapeutic protection of aRBD-2-5-Fc in hamster infected with the current circulating Omicron variant. Consistent with excellent *in vitro* neutralization activities, aRBD-2-5-Fc successfully protect the hamsters infected with Omicron at a dose of 10 mg/kg either pre- or post-challenge, there were no infectious virus detected in trachea and lungs, and virus RNA levels in these tissues were also reduced compared with control group (**Fig. 5**).

In summary, we discovered a novel conserve epitope on RBD targeted by aRBD-2, and aRBD-2 targeting this epitope is resistant to VOC escape. Hetero-bivalent tandem constructed with two Nbs with non-overlapping epitopes are beneficial for improving affinities and neutralizing activities toward VOCs. This study provides valuable insights into the development of better therapeutics for the treatment of COVID-19 caused by VOCs.

## Materials and methods

### Protein preparation

SARS-CoV-2 RBDs were prepared as our previous study^43^. Briefly, the coding sequences for SARS-CoV-2 RBD (amino acids 321 to 591) of original WT, Alpha (N501Y), Beta (K417N, E484K, N501Y), Gamma (K417T, E484K, N501Y), Delta (L452R, T478K), Delta plus (K417N, L452R, T478K), Kappa (L452R, E484Q) and Lambda (L452R, F490S) variant were cloned into pTT5 vector, which the C terminal contain a TEV cleavage site and a human IgG1 Fc. The recombinant vectors were transiently transfected into HEK293F cells with polyethyleneimine (Polyscience). After three days of expression, fusion protein was purified from cell supernatant using protein A column (GE healthcare). After digestion with TEV protease, the Fc fragment was removed by a second protein A column purification, and the TEV protease was removed by a Nickel column (GE healthcare). Omicron RBD protein was purchased from Sino Biological. Nbs, hetero-Nbs and IgG1 Fc fused hetero-Nbs including aRBD2, aRBD5, aRBD7, aRBD2-5-Fc, aRBD2-7-Fc, aRBD2-5-Fc, and aRBD2-7-Fc were prepared similarly.

### Surface Plasmon Resonance (SPR)

SPR measurements were performed at 25°C using a BIAcore T200 system. Nb-Fc was diluted to a concentration of 5 μg/ml with sodium acetate (pH 4.5) and immobilized on a CM5 chip (GE Healthcare) at a level of ∼150 response units. All proteins were exchanged into the running buffer (PBS containing 0.05% tween-20, pH 7.4), and the flow rate was 30 mL/min. The blank channel of the chip served as the negative control. For affinity measurements, a series of different concentrations of RBD flowed over the sensorchip. After each cycle, the chip was regenerated with 50 mM NaOH buffer for 60 to 120 s. The sensorgrams were fitted with a 1:1 binding model using Biacore evaluation software.

### Enzyme-linked immunosorbent assay (ELISA)

SARS-CoV-2 RBD was coated onto Immuno-MaxiSorb plates (Nunc) at final concentration of 2 μg/mL for 2 hr at 4°C. The plates were washed with PBS then blocked with MPBS (PBS containing 5% defatted milk) for 2 hr at room temperature. Serially diluted Nb-Fc solutions were added to the plates, followed by incubation for 1 hr at room temperature. After four washes with PBST (PBS containing 0.1% tween-20), bound Nb-Fc were detected with a HRP-conjugated anti-IgG1 Fc antibody (Sino Biological). After incubation for 2 hr at room temperature, the plates were washed 4 times with PBST. 100 μL per well TMB (Beyotime) was added and reacted under dark for 5 min,, 50 μL per well of H_2_SO_4_ (1 M) was added to stop the reaction. OD_450_ was read by a Synergy H1 plate reader (Biotek). The data was analyzed using GraphPad Prism software.

### Micro-neutralization assay by counting infected cells

A micro-neutralization assay by counting infected cells was employed to evaluate neutralizing activity of Nb-Fc fusions. Briefly, Nb-Fc in a 3-fold dilution concentration series were incubated with 200 PFU of SARS-CoV-2 Alpha virus (England 204820464/2020), Gamma virus (Japan TY7-503/2021) and Kappa virus (USA/CA-Stanford-15_S02/2021) for 30 min, respectively. The antibody and virus mixture was then added to Vero E6 cells in 96-well plates (Corning). After 1 h, the supernatant was removed from the wells, and the cells were washed with PBS and overlaid with Dulbecco modified Eagle medium (DMEM) containing 0.5% methylcellulose. After 2days of infection, the cells were fixed with 4% paraformaldehyde, permeabilized with 0.1% Triton-100, blocked with DMEM containing 10% fetal bovine serum, and stained with a rabbit monoclonal antibody against SARS-CoV-2 NP (GeneTex, GTX635679) and an Alexa Fluor 488-conjugated goat anti-mouse secondary antibody (Thermo Fisher Scientific). Hoechst 33342 was added in the final step to counterstain the nuclei. Fluorescence images of the entire well were acquired with a 4 X objective in a Cytation 5 (BioTek). The total number of cells indicated by the nuclei staining and the infected cells indicated by the NP staining were quantified with the cellular analysis module of the Gen5 software (BioTek). The data was analyzed using GraphPad Prism software.

### Plaque reduction neutralization (PRNT) assay

Neutralizing activity of Nb-Fc fusions was also evaluated using PRNT assay as our previous study^57^ with slight modification. Briefly, Vero E6 cells were seeded overnight in 24-well culture plates at 1.5 ×10^5^ per well. Nb-Fc were serially diluted five-fold in DMEM containing 2.5% FBS were incubated with equal volume of 75 PFU of SARS-CoV-2 WT virus (IVCAS 6.7512), Beta virus (NPRC2.062100001), Delta virus (GWHBEBW01000000) and Omicron virus (CCPM-B-V-049-2112-18) at 37°C for 1 hr, respectively. Then, the mixture was added to the cells. Cells infected with virus without antibody addition and cells without virus were used as controls. After an additional 1 hr incubation at 37°C, the antibody -virus mixture was removed, and DMEM containing 2.5% FBS and 0.9% carboxymethy lcellulose were added. Plates were fixed with 8% paraformaldehyde and stained with 0.5% crystal violet and rinsed thoroughly with water 3 days later, plaques were then enumerated and the neutralization IC_50_ was calculated using GraphPad Prism software.

### aRBD-2-5-Fc prophylactic protection in mice

Female K-18 hACE2 transgenic mice (The Jackson Laboratory) were administered with one dose of aRBD-2-5-Fc (10 mg/kg, i.p.) 24 hr before infected with 2 ×10^4^ PFU of original SARS-CoV-2 (i.n.), and female A/J mice (The Jackson Laboratory) were administered with one dose of aRBD-2-5-Fc (1 mg/kg and 0.1 mg/kg, i.p.) 24 hr before infected with 1 ×10^5^ PFU of mouse-adapted SARS-CoV-2 (intratracheally), body weight and death of the mice were monitored daily for 7 days post infection. All operations with authentic viruses were performed in the biosafety level 3 (BSL-3) facility.

### aRBD-2-5-Fc prophylactic and therapeutic protection in Syrian golden hamster

For prophylactic evaluation, female Syrian hamsters (five-to-six weeks of age) were administrated (i.p.) with one dose of 10 mg/kg of aRBD-2-5-Fc (n = 5) or PBS (n = 6) after anesthetized by chamber induction (5 L 100% O_2_/min and 3–5% isoflurane). 24 hr later, the hamsters were infected (i.n.) with 1 ×10^4^ PFU of SARS-CoV-2 Omicron virus in 100 μL of PBS. For therapeutic evaluation of aRBD-2-5-Fc, hamsters were challenged (i.n.) with 1 ×10^4^ PFU of SARS-CoV-2 Omicron virus. Three hr later, the hamsters were administrated with one dose of 10 mg/kg of aRBD-2-5-Fc (i.p.) (n = 5) or PBS (n = 6). Animals were weighed daily and euthanized at 4 d.p.i., tissues (trachea and lungs) were harvested for analysis of virus RNA copies and titers. All operations were performed in the biosafety level 3 (BSL-3) facility.

### Virus RNA copies and titers

Viral RNA in the tissue homogenates was quantified by one-step real-time RT-PCR as described before ^58^. Briefly, viral RNA was purified using the QIAamp Viral RNA Mini Kit (Qiagen), and quantified with HiScript^®^ II One Step qRT-PCR SYBR^®^ Green Kit (Vazyme Biotech Co., Ltd) with the primers ORF1ab-F(5’-CCCTGTGGGTTTTACACTTAA-3’) and ORF1ab-R (5’-ACGATTGTGCATCAGCTGA-3’). The amplification procedure was set up as: 50°C for 3 min, 95°C for 30 s followed by 40 cycles consisting of 95°C for 10 s, 60°C for 30 s.

Virus titer was determined with plaque assay as previously described with slight modification ^57^. Briefly, virus samples were serially ten-fold diluted with DMEM containing 2.5% FBS, and inoculated to Vero E6 cells cultured overnight at 1.5 × 10^5^ /well in 24-well plates; after incubating at 37°C for 1 h, the inoculate was replaced with DMEM containing 2.5% FBS and 0.9% carboxymethyl-cellulose. The plates were fixed with 8% paraformaldehyde and stained with 0.5% crystal violet 3 days later. Virus titer was calculated with the dilution gradient with 10∼100 plaques.

### Crystallization and data collection

Purified SARS-CoV-2 RBD was mixed with aRBD-5 and RBD-tir2 was mixed with aRBD-2-7 in a molar ratio of 1: 1.2 to form complexes. To remove excess Nbs, the mixtures were further purified by gel filtration. The complex protein was concentrated to 20 mg/mL for crystallization screening. Sitting-drop vapor diffusion method was applied to obtain the crystals of complexes by mixing 0.2 μl of protein complexes with an equal volume of reservoir solution. Optimized crystal of RBD in complex with aRBD5 was achieved in 0.1 M Sodium cacodylate at pH 5.5, 25% (w/v) PEG 4000 for about 1 month at 18°C, while crystal of RBD-tir2: aRBD2-7 was grown in 0.1M (NH_4_)_2_SO_4_, 0.1M Tris-HCl pH7.5, 20% (w/v) PEG1500 for about 1 month at 18°C. For data collection, single crystals were flashed-cooled in liquid nitrogen after immersing in the cryoprotectant composed of 15% (v/v) glycerol for the crystals of RBD-tr2 complexed with aRBD2-7 and 20% (v/v) ethylene glycol for the crystals of RBD in complex with aRBD-5 in the containing reservoir solution for few seconds. Diffraction data were collected at Shanghai Synchrotron Radiation Facility (SSRF) beamline BL19U1 for RBD in complex with aRBD5 at the wavelength of 0.97852 Å and BL02U1 for RBD-tr2 complexed with aRBD2-7 at the wavelength of 0.97918 Å, respectively.

### Structural determination

Data were processed with XDS^59^. Initial phases were solved by molecular replacement method with Phaser ^60^ from the CCP4i program package ^61^, using SARS-CoV-2 RBD/ACE2 (PDB ID: 6M0J) and RSV/F-VHH-4 (PDB ID: 5TP3) as search models for aRBD-5 in complex with RBD, and SARS-CoV-2 RBD/ACE2 (PDB ID: 6M0J), TcdB-B1/B39 VHH (PDB ID:4NC2) and Vsig4/Nb119 (PDB ID: 5IMK) were used as search models for RBD-tr2 complexed with: aRBD2-7. Subsequent models building and refinement were achieved using COOT and Phenix ^62^. All structural figures were prepared by PyMOL.

### Statistical analysis

All statistical analyses were performed in GraphPad Prism. An unpaired t-test with a Welch’s correction for unequal standard deviations was used for comparisons of two groups. The asterisks shown in the figures refer to the level of significance: *p ≤ 0.05; **p ≤ 0.01.

## Acknowledgement

We thank all of the staff who participated in this work for their important contributions. We thank Jia Wu, Jun Liu and Hao Tang from Wuhan Institute of Virology for the management of BSL-3 facility. We also thank National Virus Resource Center for providing Omicron variant (CCPM-B-V-049-2112-18). This study was supported by the Strategic Priority Research Program of the Chinese Academy of Sciences (Grant No. XDB29030104), the National Natural Science Foundation of China (Grant No. 31870731, 32100745), the Fundamental Research Funds for the Central Universities (Grant No. WK9100000019), the Fundamental Research Funds for the Central Universities (Grant No. WK100000019).

## Author Contributions

T.J. S.C. and Y.X. provided funding, designed the study, participated in data analysis, and revised the manuscript. H.M. X.Z. P.Z. P.D. W.Z. S.C. and H.Z. designed the study, performed experiments, analyzed the data, and drafted the manuscript. Other authors participated in the experiments and/or revised the manuscript.

## Supplementary information

### Supplementary Figures

**Supplementary Fig. 1.**
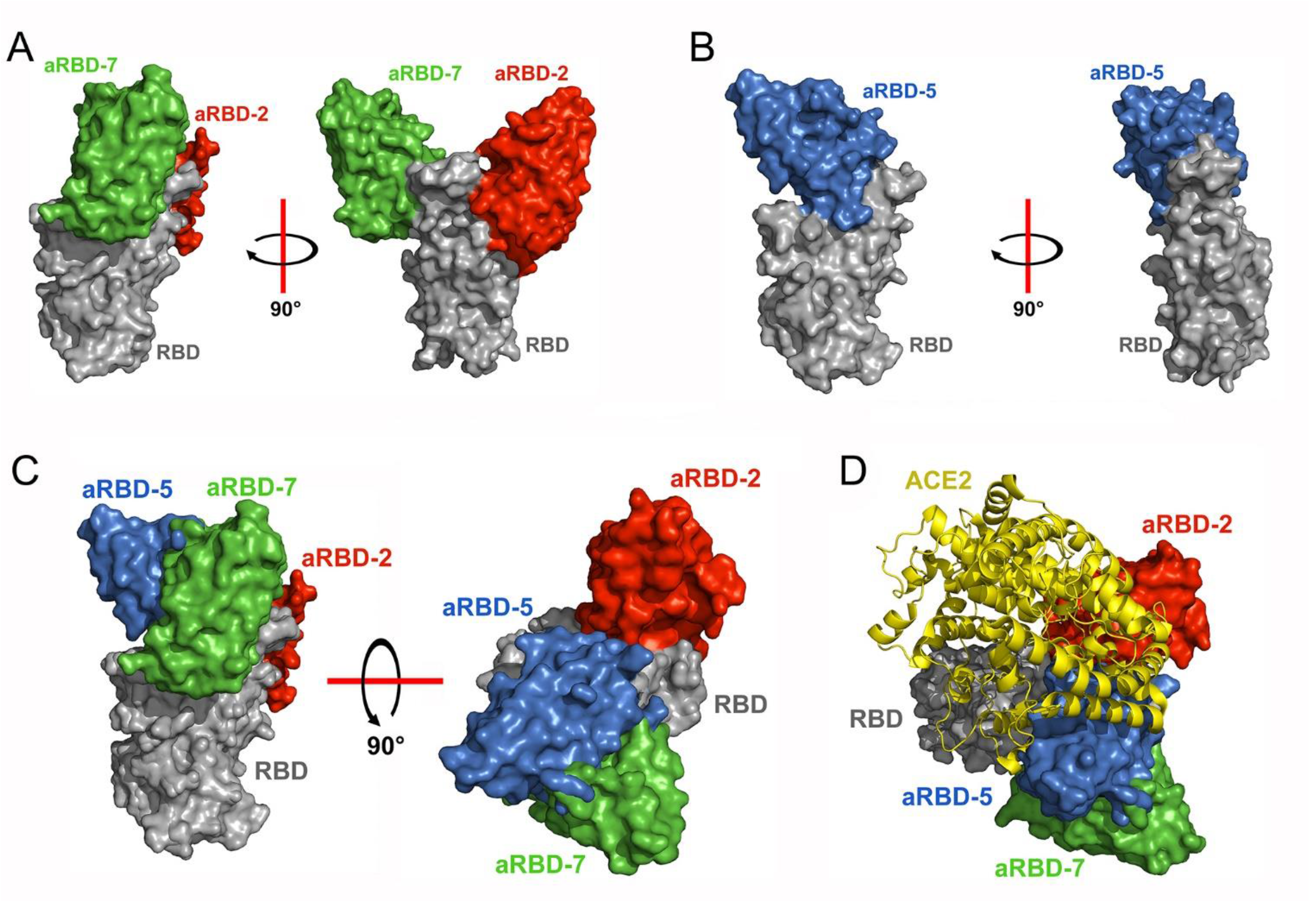
Crystal structures of aRBD-2, aRBD-5 and aRBD-7 in complex with RBD using surface presentation. (A) and (B) is the overall structure of aRBD-2 (red) and aRBD-7 (green) bound to RBD (grey) and aRBD-5 (marine) bound to RBD (grey), respectively. Overall structure of aRBD-2, aRBD-5 and aRBD-7 in complex with the RBD were aligned with each other (C) and the structure of ACE2:RBD complex (PDB: 6M0J) (D).

**Supplementary Fig. 2.**
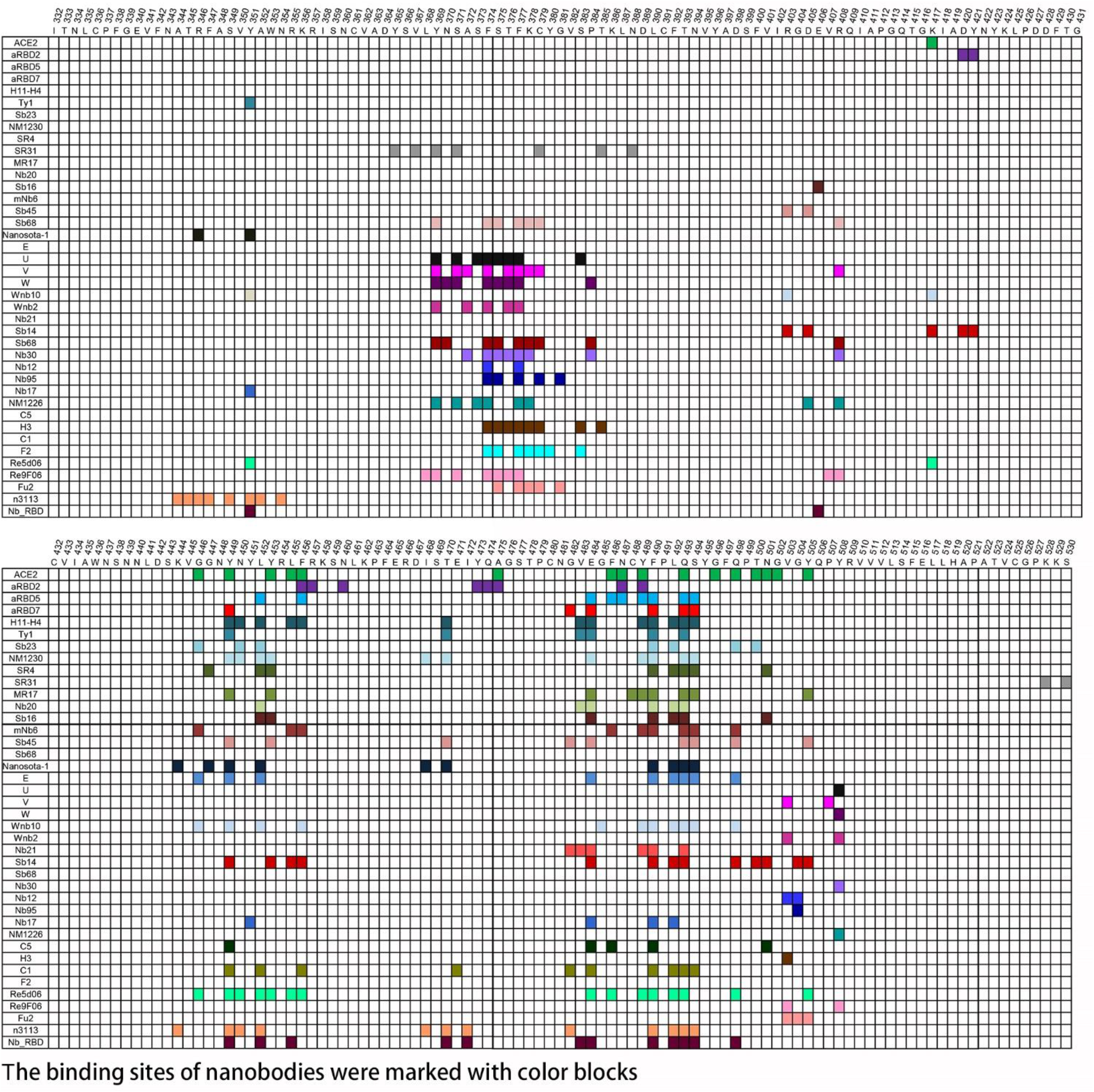
The epitopes of aRBD-2, aRBD-5 and aRBD-7 and other published nanobodies.

**Supplementary Fig. 3.**
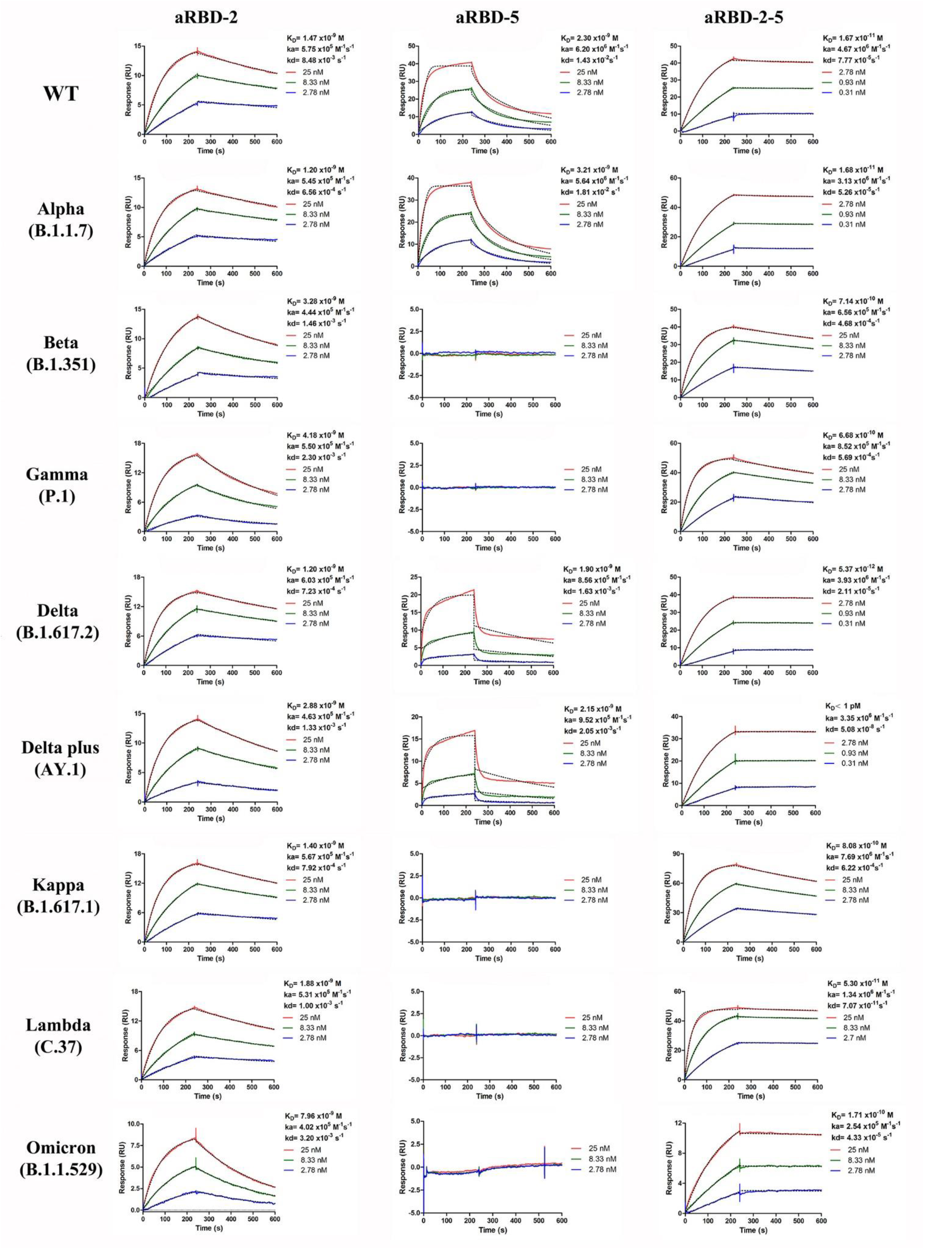
Binding affinity characterization of aRBD-2, aRBD-5 and aRBD-2-5 to the RBD of variants using SPR. Binding kinetics of original, Alpha, Beta, Gamma, Delta, Delta plus, Kappa, Lambda and Omicron variant RBD to aRBD-2, aRBD-5 and aRBD-2-5 was measured by SPR, respectively. The Nb-Fc fusions were immobilized onto a CM5 sensor chip, respectively, mutant RBDs with serially 1:3 dilutions was injected successively and monitored by the Biacore T200 system. The actual responses (colored lines) and the data fitted to a 1:1 binding model (black dotted lines) are shown.

**Supplementary Fig. 4.**
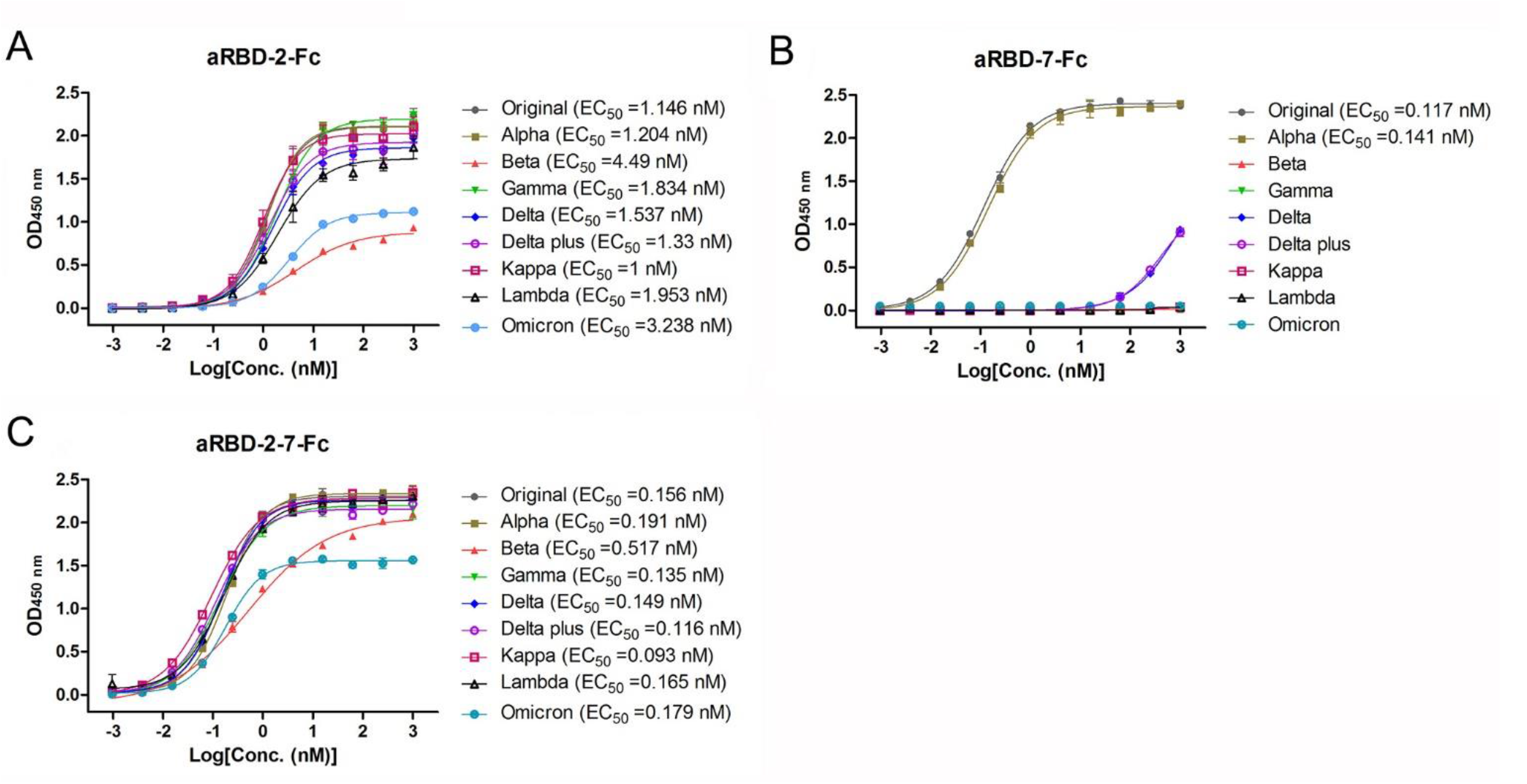
Characterization the binding of aRBD-2-Fc, aRBD-7-Fc and aRBD-2-7-Fc to the RBD of variants using ELISA. (A), (B) and (C) is the binding results of aRBD-2-Fc, aRBD-7-Fc and aRBD-2-7-Fc to the RBD of SARS-CoV-2 variants, respectively. EC_50_ was calculated by fitting the OD_450_ from serially diluted antibody with a sigmoidal dose-response curve. Error bars indicate mean ±SD from two replicates.

**Supplementary Fig. 5.**
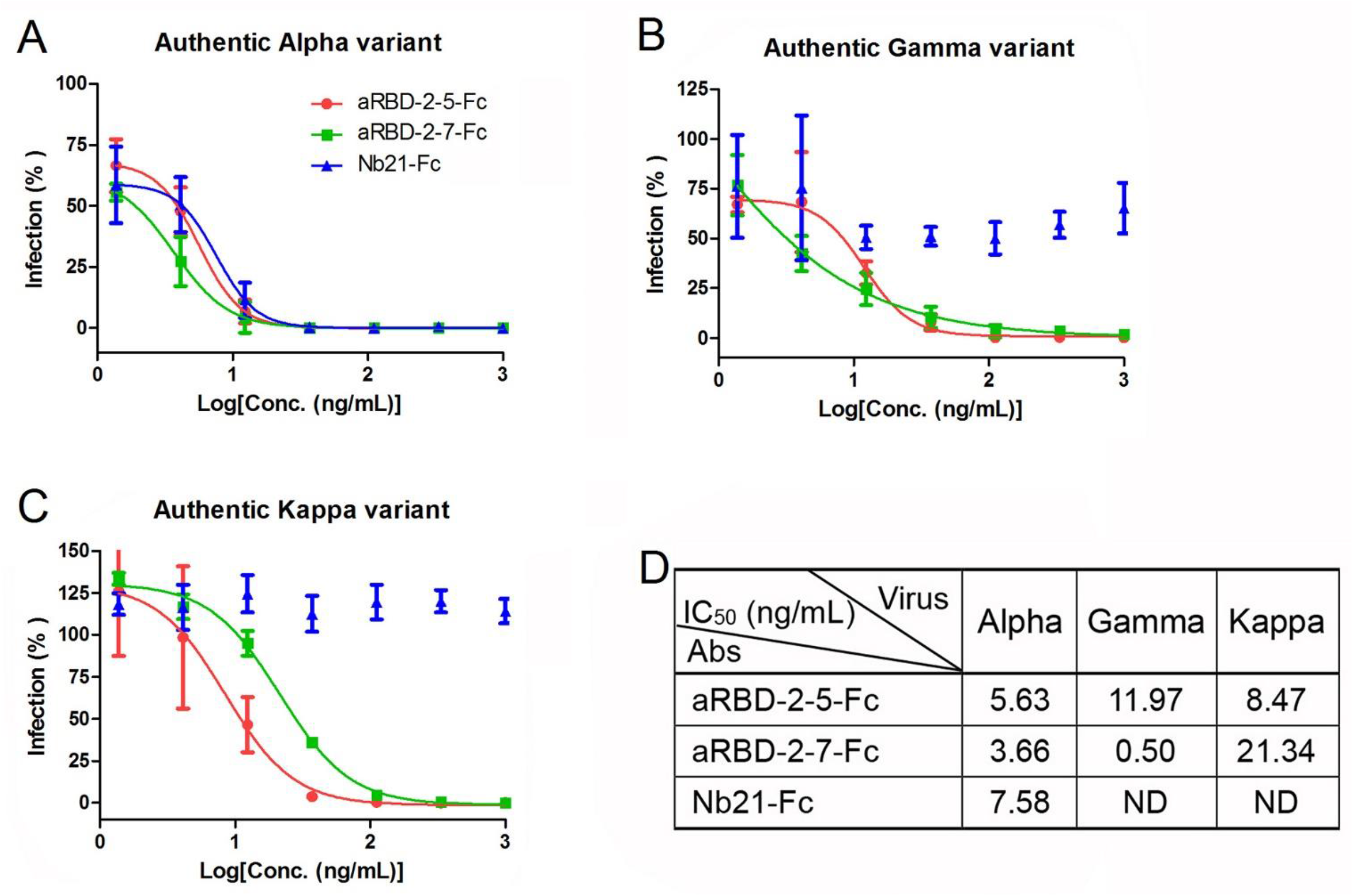
*In vitro* neutralization to SARS-CoV-2 variants by aRBD-2-5-Fc and aRBD-2-7-Fc using micro-neutralization assay. The serially diluted aRBD-2-5-Fc, aRBD-2-7-Fc and control antibody (Nb21-Fc) were incubated with ∼200 PFU of authentic Alpha (A), Gamma (B), and Kappa (C) variant. The mixture was then added to Vero E6 cells in 96-well plates. After 2 days of infection, the infected virus was stained green with a monoclonal antibody against SARS-CoV-2 NP and an Alexa Fluor 488-conjugated goat anti-mouse secondary antibody. The nucleus was stained blue with Hoechst 33342. Each experiment was performed in duplicate. The data was calculated by fitting the inhibition from serially diluted antibody to a sigmoidal dose-response curve. Error bars indicate the mean ± SD from three independent experiments. (D) is the summary of neutralizing IC_50_. *ND means neutralizing activity not detected.

**Supplementary Fig. 6.**
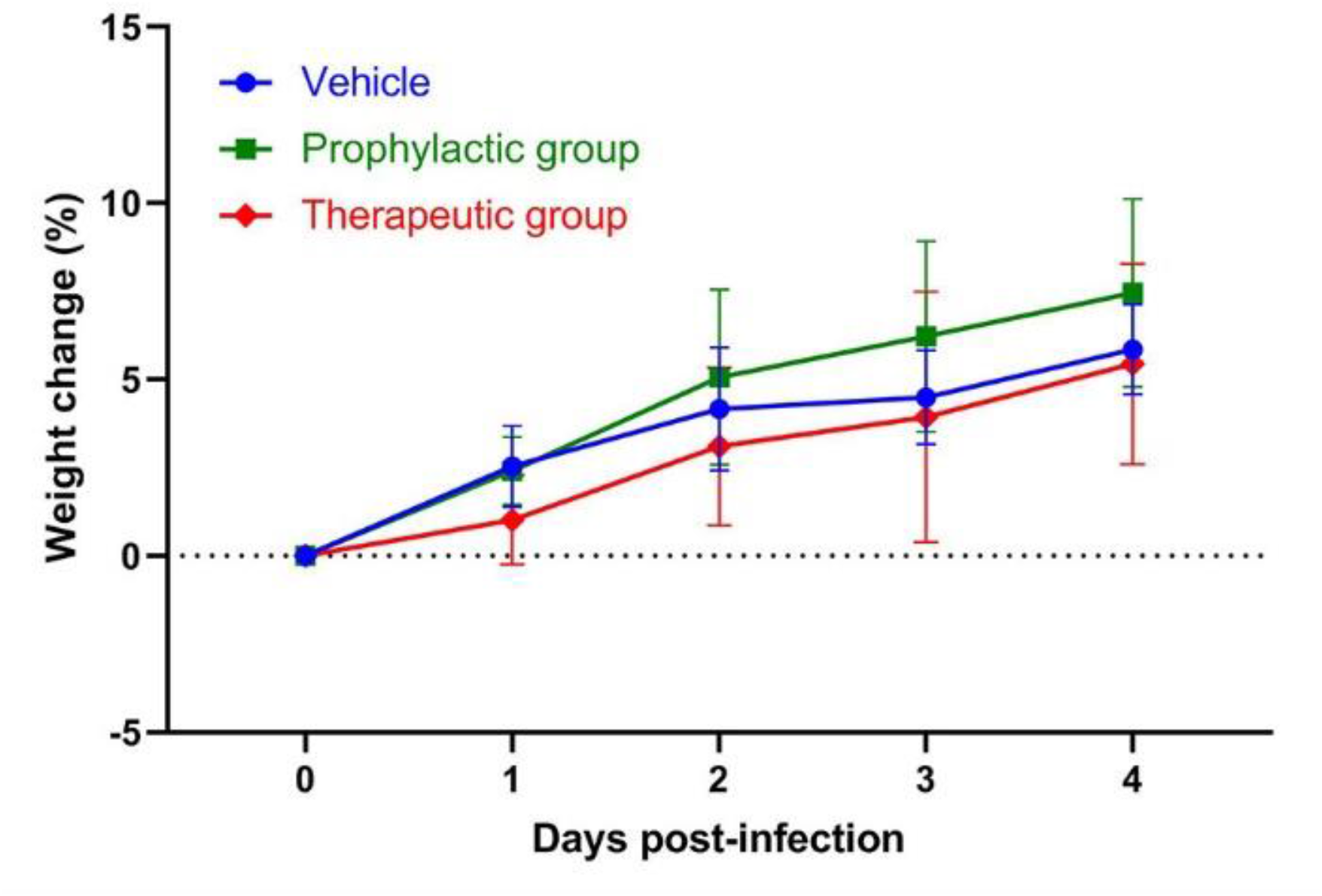
Weight change of the hamsters in prophylactic group and therapeutic group.

**Supplementary Fig. 7.**
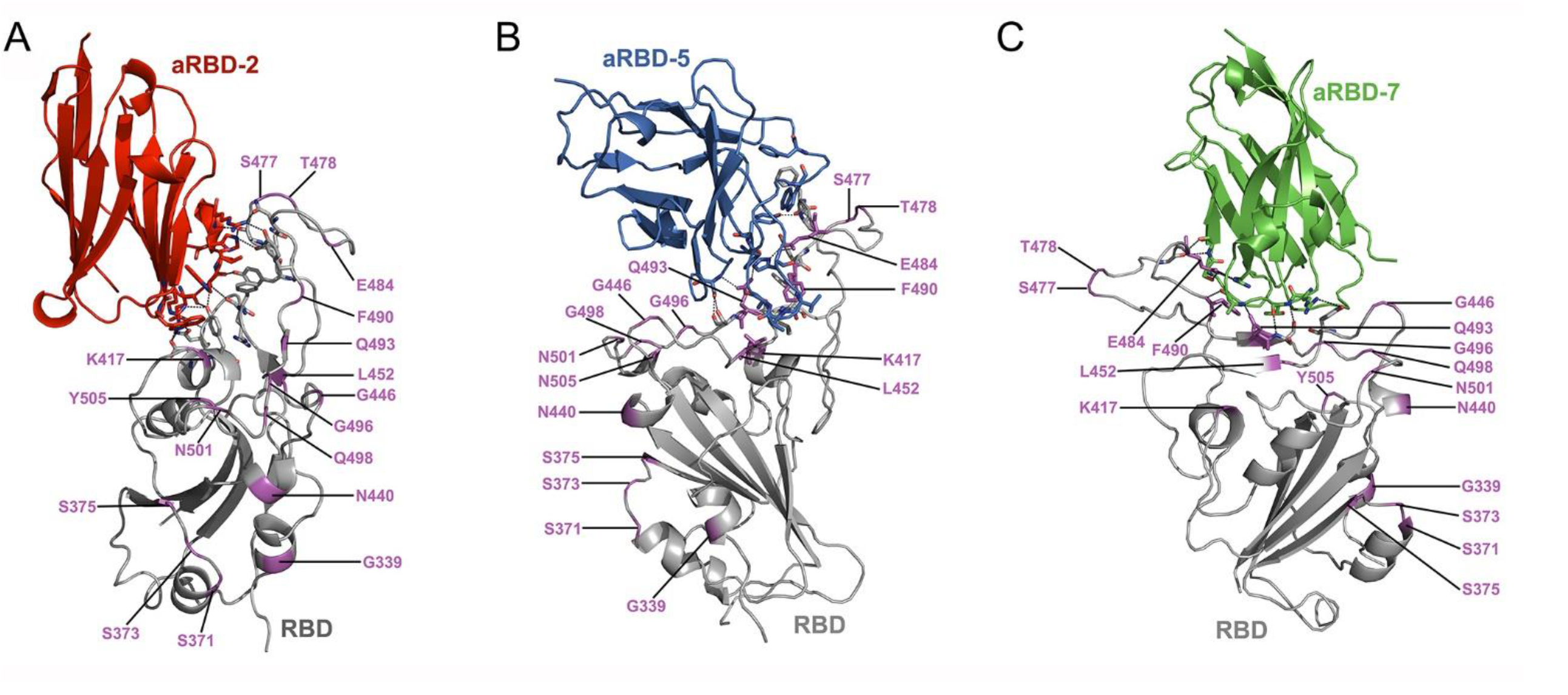
Display of mutation sites of SARS-CoV-2 variants on structures of the Nb:RBD complex. The 17 mutation sites (magenta) including G339, S371, S373, S375, K417, N440, G446, L452, S477, T478, E484, F490, Q493, G496, Q498, N501 and Y505 derived from the RBD of Alpha, Beta, Gamma, Delta, Delta plus, Kappa, Lambda and Omicron variant are displayed on the structure of aRBD-2:RBD (A), aRBD-5:RBD (B) and aRBD-7:RBD (C) complex. Residues that form interactions are shown as sticks, hydrogen bonds and salt bridges between Nbs and the RBD are shown as black dotted lines.

**Supplementary Fig. 8.**
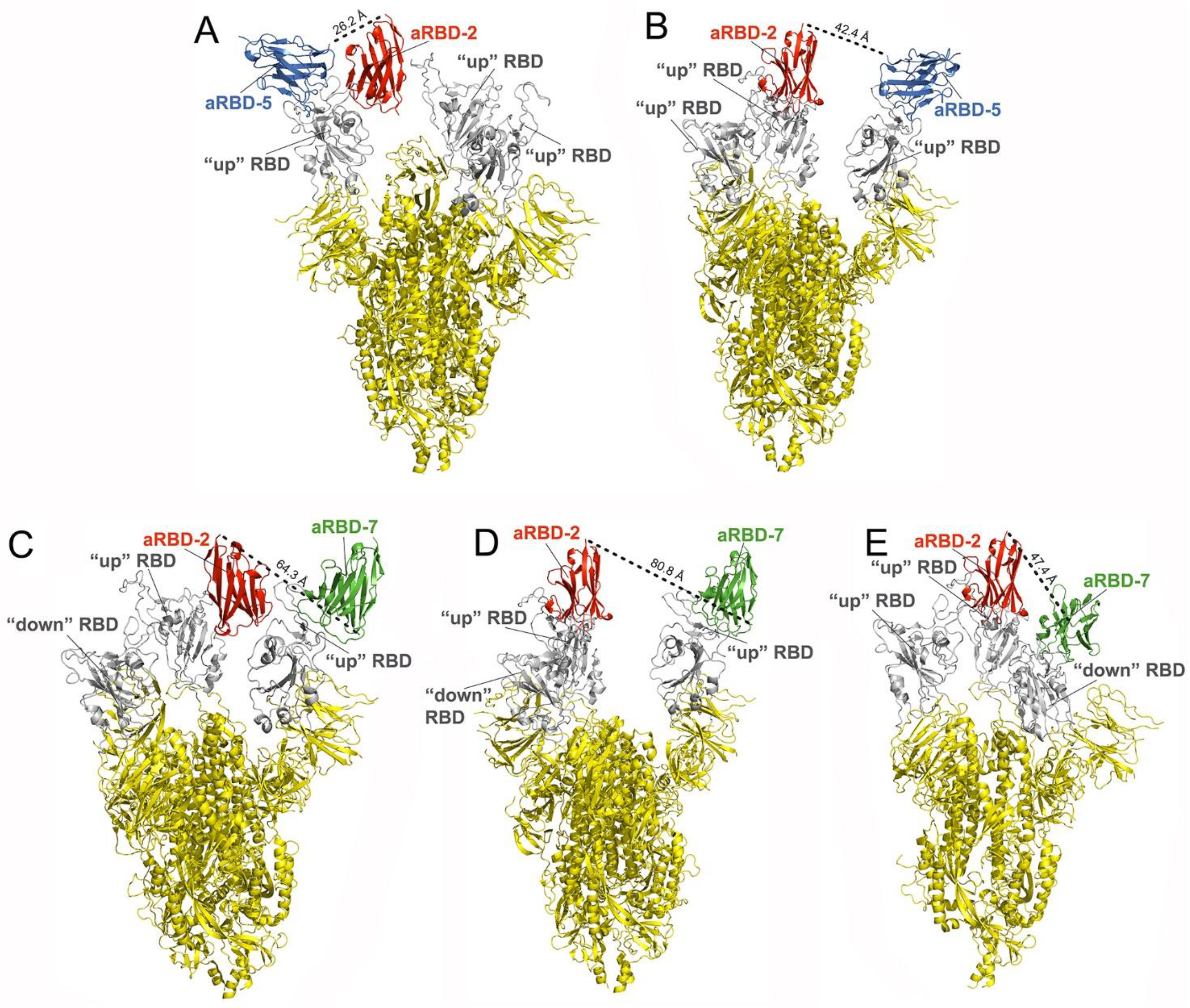
The distance between the C-terminus of aRBD-2 and the N-terminus of aRBD-5 or aRBD-7 in different binding modes. aRBD-2 and aRBD-5 in complex with RBD was aligned to the Cryo-EM structure of the trimer spike with all “up” conformation (PDB: 7KMS), and the distance between the C-terminus of aRBD-2 and the N-terminus of aRBD-5 on the same RBD (A) or on two different RBD (B) was measured. aRBD-2 and aRBD-7 in complex with RBD was aligned to the cryo-EM structure of the trimer spike with two “up” and one “down” conformation (PDB: 7KMZ), and the distance between the C-terminus of aRBD-2 and the N-terminus of aRBD-5 on the same RBD (C) or on two different “up” RBD (D) or one on “up” another on “down” RBD (E) was measured, respectively.

### Supplementary tables

**Supplementary table 1.**
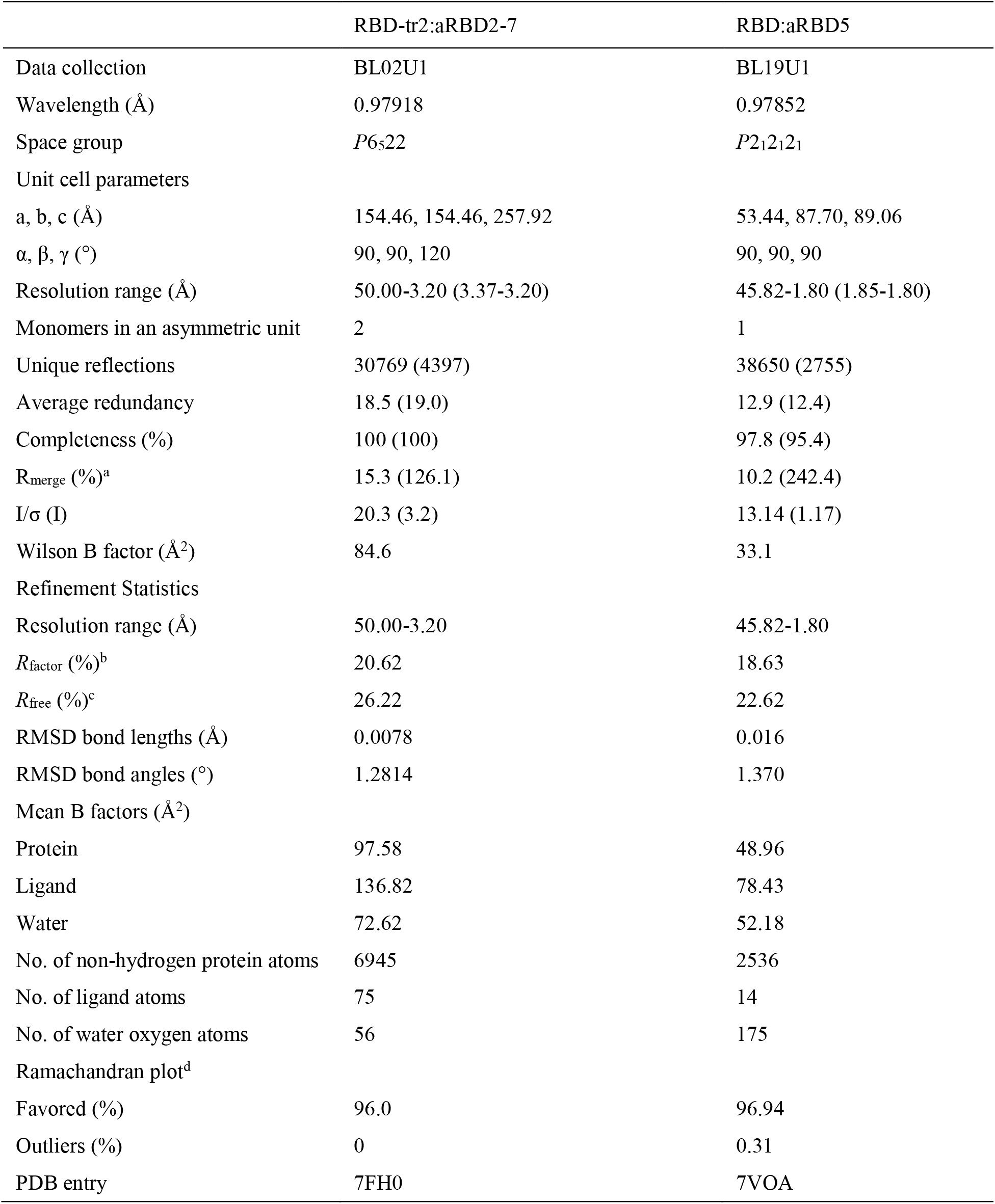
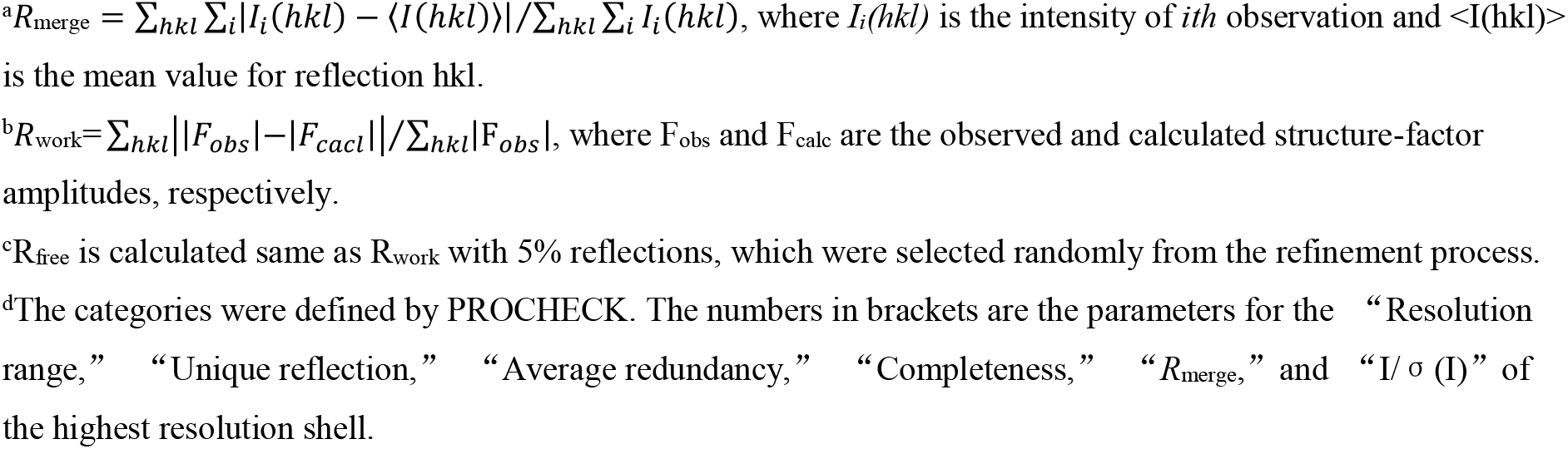
Data collection and refinement statistics

|  | RBD-tr2:aRBD2-7 | RBD:aRBD5 |
| --- | --- | --- |
| Data collection | BL02U1 | BL19U1 |
| Wavelength (Å) | 0.97918 | 0.97852 |
| Space group | *P*6<sub>5</sub>22 | *P*2<sub>1</sub>2<sub>1</sub>2<sub>1</sub> |
| Unit cell parameters |  |  |
| a, b, c (Å) | 154.46, 154.46, 257.92 | 53.44, 87.70, 89.06 |
| $\alpha$, $\beta$, $\gamma$ (°) | 90, 90, 120 | 90, 90, 90 |
| Resolution range (Å) | 50.00-3.20 (3.37-3.20) | 45.82-1.80 (1.85-1.80) |
| Monomers in an asymmetric unit | 2 | 1 |
| Unique reflections | 30769 (4397) | 38650 (2755) |
| Average redundancy | 18.5 (19.0) | 12.9 (12.4) |
| Completeness (%) | 100 (100) | 97.8 (95.4) |
| R<sub>merge</sub> (%)<sup>a</sup> | 15.3 (126.1) | 10.2 (242.4) |
| I/$\sigma$ (I) | 20.3 (3.2) | 13.14 (1.17) |
| Wilson B factor (Å<sup>2</sup>) | 84.6 | 33.1 |
| Refinement Statistics |  |  |
| Resolution range (Å) | 50.00-3.20 | 45.82-1.80 |
| R<sub>factor</sub> (%)<sup>b</sup> | 20.62 | 18.63 |
| R<sub>free</sub> (%)<sup>c</sup> | 26.22 | 22.62 |
| RMSD bond lengths (Å) | 0.0078 | 0.016 |
| RMSD bond angles (°) | 1.2814 | 1.370 |
| Mean B factors (Å<sup>2</sup>) |  |  |
| Protein | 97.58 | 48.96 |
| Ligand | 136.82 | 78.43 |
| Water | 72.62 | 52.18 |
| No. of non-hydrogen protein atoms | 6945 | 2536 |
| No. of ligand atoms | 75 | 14 |
| No. of water oxygen atoms | 56 | 175 |
| Ramachandran plot<sup>d</sup> |  |  |
| Favored (%) | 96.0 | 96.94 |
| Outliers (%) | 0 | 0.31 |
| PDB entry | 7FH0 | 7VOA |

<sup>a</sup> $R_{\text{merge}} = \sum_{hkl} \sum_i |I_i(hkl) - \langle I(hkl) \rangle| / \sum_{hkl} \sum_i I_i(hkl)$ , where $I_i(hkl)$ is the intensity of $i$ th observation and $\langle I(hkl) \rangle$ is the mean value for reflection $hkl$ .
<sup>b</sup> $R_{\text{work}} = \sum_{hkl} ||F_{\text{obs}}| - |F_{\text{calc}}|| / \sum_{hkl} |F_{\text{obs}}|$ , where $F_{\text{obs}}$ and $F_{\text{calc}}$ are the observed and calculated structure-factor amplitudes, respectively.
<sup>c</sup> $R_{\text{free}}$ is calculated same as $R_{\text{work}}$ with 5% reflections, which were selected randomly from the refinement process.
<sup>d</sup>The categories were defined by PROCHECK. The numbers in brackets are the parameters for the “Resolution range,” “Unique reflection,” “Average redundancy,” “Completeness,” “ $R_{\text{merge}}$ ,” and “ $I/\sigma(I)$ ” of the highest resolution shell.

**Supplementary table 2.** Interaction analysis results of aRBD-2, aRBD-5 and aRBD-7 binding to SARS-CoV-2 RBD.

| RBD |  | aRBD-2 |  | Bond |
| --- | --- | --- | --- | --- |
| Residue | atom | Residue | atom |  |
| D420 | OD2 | R49 | NH1 | Hydrogen bond |
| D420 | OD2 | W50 | NE1 | Hydrogen bond |
| Y421 | OH | R49 | NH1 | Salt bridge |
| Y421 | OH | W97 | N | Hydrogen bond |
| Y421 | OH | E95 | O | Hydrogen bond |
| Y421 | | R49 | | Cation-π |
| R457 | O | W97 | N | Hydrogen bond |
| N460 | N | E95 | OE1 | Hydrogen bond |
| N460 | OD1 | R49 | NH2 | Hydrogen bond |
| Y473 | OH | L98 | O | Hydrogen bond |
| Q474 | O | H104 | NE2 | Hydrogen bond |
| A475 | O | R100 | N | Hydrogen bond |
| A475 | O | H104 | NE2 | Hydrogen bond |
| N487 | OD1 | R100 | NH1 | Hydrogen bond |
| Y489 | OH | R100 | NH1 | Hydrogen bond |
| F456/Y489 |  | L98 |  | Hydrophobic interaction |

| RBD |  | aRBD-7 |  | Bond |
| --- | --- | --- | --- | --- |
| Residue | atom | Residue | atom |  |
| Y449 | OH | R101 | NH1 | Salt bridge |
| G482 | O | T58 | N | Hydrogen bond |
| E484 | OE1 | R52 | NH2 | Hydrogen bond |
| E484 | OE2 | R52 | NE | Hydrogen bond |
| E484 | OE2 | R52 | NH2 | Salt bridge |
| E484 | OE1 | S57 | OG | Hydrogen bond |
| E484 | N | T58 | O | Hydrogen bond |
| F490 | | R52 | | Cation-π |
| F490 | N | A104 | O | Hydrogen bond |
| Q493 | OE1 | A104 | N | Hydrogen bond |
| S494 | OG | T102 | N | Hydrogen bond |
| S494 | N | T102 | O | Hydrogen bond |

| RBD |  | aRBD-5 |  | Bond |
| --- | --- | --- | --- | --- |
| Residue | atom | Residue | atom |  |
| F456,Y489 |  | Y31 |  | Hydrophobic interaction |
| E484 | OE1 | H101 | N | Hydrogen bond |
| E484 | OE2 | H101 | ND1 | Hydrogen bond |
| F486 |  | V2, W112,F115 |  | Hydrophobic interaction |
| N487 | N | Y32 | OH | Hydrogen bond |
| Q493 | OE1 | S54 | N | Hydrogen bond |
| Q493 | NE2 | H101 | NE2 | Hydrogen bond |
| S494 | O | S54 | OG | Hydrogen bond |
| S494 | OG | S54 | OG | Hydrogen bond |
| L452, F490 and<br>L492 |  | V103, V104 and<br>A105 |  | Hydrophobic interaction |

## References

1 Singh D, Yi SV. On the origin and evolution of SARS-CoV-2. Experimental & Molecular Medicine 2021; 53:537–547.

2 Rochman ND, Wolf YI, Faure G, Mutz P, Zhang F, Koonin EV. Ongoing global and regional adaptive evolution of SARS-CoV-2. Proceedings of the National Academy of Sciences of the United States of America 2021; 118.

3 Li J, Lai S, Gao GF, Shi W. The emergence, genomic diversity and global spread of SARS-CoV-2. Nature 2021; 600:408–418.

4 Tao K, Tzou PL, Nouhin J et al. The biological and clinical significance of emerging SARS-CoV-2 variants. Nature reviews Genetics 2021:1–17.

5 Poudel S, Ishak A, Perez-Fernandez J et al. Highly mutated SARS-CoV-2 Omicron variant sparks significant concern among global experts - What is known so far? Travel medicine and infectious disease 2022; 45:102234.

6 Cele S, Jackson L, Khoury DS et al. Omicron extensively but incompletely escapes Pfizer BNT162b2 neutralization. Nature 2021.

7 Lu L, Mok BW-Y, Chen L et al. Neutralization of SARS-CoV-2 Omicron variant by sera from BNT162b2 or Coronavac vaccine recipients. medRxiv : the preprint server for health sciences 2021.

8 COVID-19 Vaccine Breakthrough Infections Reported to CDC - United States, January 1-April 30, 2021. MMWR Morbidity and mortality weekly report 2021; 70:792–793.

9 Maxmen A. The fight to manufacture COVID vaccines in lower-income countries. Nature 2021; 597:455–457.

10 DeFrancesco L. Whither COVID-19 vaccines? Nature biotechnology 2020; 38:1132–1145.

11 Zohar T, Alter G. Dissecting antibody-mediated protection against SARS-CoV-2. Nature reviews Immunology 2020; 20:392–394.

12 Taylor PC, Adams AC, Hufford MM, de la Torre I, Winthrop K, Gottlieb RL. Neutralizing monoclonal antibodies for treatment of COVID-19. Nature reviews Immunology 2021; 21:382–393.

13 Planas D, Saunders N, Maes P et al. Considerable escape of SARS-CoV-2 Omicron to antibody neutralization. Nature 2021.

14 Dejnirattisai W, Huo J, Zhou D et al. SARS-CoV-2 Omicron-B.1.1.529 leads to widespread escape from neutralizing antibody responses. Cell 2022; 185:467–484 e415.

15 VanBlargan LA, Errico JM, Halfmann PJ et al. An infectious SARS-CoV-2 B.1.1.529 Omicron virus escapes neutralization by therapeutic monoclonal antibodies. Nature medicine 2022:1–6.

16 Cao Y, Wang J, Jian F et al. Omicron escapes the majority of existing SARS-CoV-2 neutralizing antibodies. Nature 2021.

17 Saelens X, Schepens B. Single-domain antibodies make a difference. Science (New York, NY) 2021; 371:681–682.

18 Sasisekharan R. Preparing for the Future - Nanobodies for Covid-19? The New England journal of medicine 2021; 384:1568–1571.

19 Czajka TF, Vance DJ, Mantis NJ. Slaying SARS-CoV-2 One (Single-domain) Antibody at a Time. Trends in microbiology 2021; 29:195–203.

20 Ahmad J, Jiang J, Boyd LF et al. Structures of synthetic nanobody-SARS-CoV-2 receptor-binding domain complexes reveal distinct sites of interaction. The Journal of biological chemistry 2021; 297:101202.

21 Custódio TF, Das H, Sheward DJ et al. Selection, biophysical and structural analysis of synthetic nanobodies that effectively neutralize SARS-CoV-2. Nature communications 2020; 11:5588.

22 Huo J, Le Bas A, Ruza RR et al. Neutralizing nanobodies bind SARS-CoV-2 spike RBD and block interaction with ACE2. Nature structural & molecular biology 2020; 27:846–854.

23 Hanke L, Vidakovics Perez L, Sheward DJ et al. An alpaca nanobody neutralizes SARS-CoV-2 by blocking receptor interaction. Nature communications 2020; 11:4420.

24 Yao H, Cai H, Li T et al. A high-affinity RBD-targeting nanobody improves fusion partner’s potency against SARS-CoV-2. PLoS pathogens 2021; 17:e1009328.

25 Schoof M, Faust B, Saunders RA et al. An ultrapotent synthetic nanobody neutralizes SARS-CoV-2 by stabilizing inactive Spike. Science (New York, NY) 2020; 370:1473–1479.

26 Koenig PA, Das H, Liu H et al. Structure-guided multivalent nanobodies block SARS-CoV-2 infection and suppress mutational escape. Science (New York, NY) 2021; 371.

27 Wagner TR, Ostertag E, Kaiser PD et al. NeutrobodyPlex-monitoring SARS-CoV-2 neutralizing immune responses using nanobodies. EMBO reports 2021; 22:e52325.

28 Pymm P, Adair A, Chan LJ et al. Nanobody cocktails potently neutralize SARS-CoV-2 D614G N501Y variant and protect mice. Proceedings of the National Academy of Sciences of the United States of America 2021; 118.

29 Xu J, Xu K, Jung S et al. Nanobodies from camelid mice and llamas neutralize SARS-CoV-2 variants. Nature 2021; 595:278–282.

30 Sun D, Sang Z, Kim YJ et al. Potent neutralizing nanobodies resist convergent circulating variants of SARS-CoV-2 by targeting diverse and conserved epitopes. Nature communications 2021; 12:4676.

31 Huo J, Mikolajek H, Le Bas A et al. A potent SARS-CoV-2 neutralising nanobody shows therapeutic efficacy in the Syrian golden hamster model of COVID-19. Nature communications 2021; 12:5469.

32 Güttler T, Aksu M, Dickmanns A et al. Neutralization of SARS-CoV-2 by highly potent, hyperthermostable, and mutation-tolerant nanobodies. The EMBO journal 2021; 40:e107985.

33 Li T, Cai H, Yao H et al. A synthetic nanobody targeting RBD protects hamsters from SARS-CoV-2 infection. Nature communications 2021; 12:4635.

34 Lu Q, Zhang Z, Li H et al. Development of multivalent nanobodies blocking SARS-CoV-2 infection by targeting RBD of spike protein. Journal of nanobiotechnology 2021; 19:33.

35 Chi X, Liu X, Wang C et al. Humanized single domain antibodies neutralize SARS-CoV-2 by targeting the spike receptor binding domain. Nature communications 2020; 11:4528.

36 Gai J, Ma L, Li G et al. A potent neutralizing nanobody against SARS-CoV-2 with inhaled delivery potential. MedComm 2021; 2:101–113.

37 Xiang Y, Nambulli S, Xiao Z et al. Versatile and multivalent nanobodies efficiently neutralize SARS-CoV-2. Science (New York, NY) 2020; 370:1479–1484.

38 Wrapp D, De Vlieger D, Corbett KS et al. Structural Basis for Potent Neutralization of Betacoronaviruses by Single-Domain Camelid Antibodies. Cell 2020; 181:1436–1441.

39 Hanke L, Das H, Sheward DJ et al. A bispecific monomeric nanobody induces spike trimer dimers and neutralizes SARS-CoV-2 in vivo. Nature communications 2022; 13:155.

40 Chi X, Zhang X, Pan S et al. An ultrapotent RBD-targeted biparatopic nanobody neutralizes broad SARS-CoV-2 variants. Signal transduction and targeted therapy 2022; 7:44.

41 Yang Z, Wang Y, Jin Y et al. A non-ACE2 competing human single-domain antibody confers broad neutralization against SARS-CoV-2 and circulating variants. Signal transduction and targeted therapy 2021; 6:378.

42 De Meyer T, Muyldermans S, Depicker A. Nanobody-based products as research and diagnostic tools. Trends in biotechnology 2014; 32:263–270.

43 Ma H, Zeng W, Meng X et al. Potent Neutralization of SARS-CoV-2 by Hetero-bivalent Alpaca Nanobodies Targeting the Spike Receptor-Binding Domain. Journal of virology 2021; 95.

44 Pinto D, Park YJ, Beltramello M et al. Cross-neutralization of SARS-CoV-2 by a human monoclonal SARS-CoV antibody. Nature 2020; 583:290–295.

45 Leist SR, Dinnon KH, 3rd, Schäfer A et al. A Mouse-Adapted SARS-CoV-2 Induces Acute Lung Injury and Mortality in Standard Laboratory Mice. Cell 2020; 183:1070–1085 e1012.

46 Halfmann PJ, Iida S, Iwatsuki-Horimoto K et al. SARS-CoV-2 Omicron virus causes attenuated disease in mice and hamsters. Nature 2022.

47 Guan Y, Zheng BJ, He YQ et al. Isolation and characterization of viruses related to the SARS coronavirus from animals in southern China. Science (New York, NY) 2003; 302:276–278.

48 Li W, Shi Z, Yu M et al. Bats are natural reservoirs of SARS-like coronaviruses. Science (New York, NY) 2005; 310:676–679.

49 Ge XY, Li JL, Yang XL et al. Isolation and characterization of a bat SARS-like coronavirus that uses the ACE2 receptor. Nature 2013; 503:535–538.

50 Yang XL, Hu B, Wang B et al. Isolation and Characterization of a Novel Bat Coronavirus Closely Related to the Direct Progenitor of Severe Acute Respiratory Syndrome Coronavirus. Journal of virology 2015; 90:3253–3256.

51 Hu B, Zeng LP, Yang XL et al. Discovery of a rich gene pool of bat SARS-related coronaviruses provides new insights into the origin of SARS coronavirus. PLoS pathogens 2017; 13:e1006698.

52 Xiao K, Zhai J, Feng Y et al. Isolation of SARS-CoV-2-related coronavirus from Malayan pangolins. Nature 2020; 583:286–289.

53 Zhou H, Ji J, Chen X et al. Identification of novel bat coronaviruses sheds light on the evolutionary origins of SARS-CoV-2 and related viruses. Cell 2021; 184:4380–4391 e4314.

54 Lan J, Ge J, Yu J et al. Structure of the SARS-CoV-2 spike receptor-binding domain bound to the ACE2 receptor. Nature 2020; 581:215–220.

55 Greaney AJ, Loes AN, Crawford KHD et al. Comprehensive mapping of mutations in the SARS-CoV-2 receptor-binding domain that affect recognition by polyclonal human plasma antibodies. Cell host & microbe 2021; 29:463–476 e466.

56 Wang P, Nair MS, Liu L et al. Antibody resistance of SARS-CoV-2 variants B.1.351 and B.1.1.7. Nature 2021; 593:130–135.

57 Zhang Q, Zhang H, Gao J et al. A serological survey of SARS-CoV-2 in cat in Wuhan. Emerging microbes & infections 2020; 9:2013–2019.

58 Feng L, Wang Q, Shan C et al. An adenovirus-vectored COVID-19 vaccine confers protection from SARS-COV-2 challenge in rhesus macaques. Nature communications 2020; 11:1–11.

59 Kabsch W. XDS. Acta crystallographica Section D, Biological crystallography 2010; 66:125–132.

60 McCoy AJ, Grosse-Kunstleve RW, Adams PD, Winn MD, Storoni LC, Read RJ. Phaser crystallographic software. Journal of applied crystallography 2007; 40:658–674.

61 Potterton E, Briggs P, Turkenburg M, Dodson E. A graphical user interface to the CCP4 program suite. Acta crystallographica Section D, Biological crystallography 2003; 59:1131–1137.

62 Adams PD, Afonine PV, Bunkóczi G et al. PHENIX: a comprehensive Python-based system for macromolecular structure solution. Acta crystallographica Section D, Biological crystallography 2010; 66:213–221.

